# Canonical Wnt transcriptional complexes are essential for induction of nephrogenesis but not maintenance or proliferation of nephron progenitors

**DOI:** 10.1101/2023.08.20.554044

**Authors:** Helena Bugacov, Balint Der, Sunghyun Kim, Nils O. Lindström, Andrew P. McMahon

**Author notes:** Corresponding author: Dr. Andrew P. McMahon.

## Abstract

Wnt regulated transcriptional programs are associated with both the maintenance of mammalian nephron progenitor cells (NPC) and their induction, initiating the process of nephrogenesis. How opposing transcriptional roles are regulated remain unclear. Using an *in vitro* model replicating *in vivo* events, we examined the requirement for canonical Wnt transcriptional complexes in NPC regulation. In canonical transcription, Lef/Tcf DNA binding proteins associate the transcriptional co-activator β-catenin. Wnt signaling is readily substituted by CHIR99021, a small molecule antagonist of glycogen synthase kinase-3β (GSK3β). GSK3β inhibition blocks Gskβ-dependent turnover of β-catenin, enabling formation of Lef/Tcf/β-catenin transcriptional complexes, and enhancer-mediated transcriptional activation. Removal of β-catenin activity from NPCs under cell expansion conditions (low CHIR) demonstrated a non-transcriptional role for β-catenin in the CHIR-dependent proliferation of NPCs. In contrast, CHIR-mediated induction of nephrogenesis, on switching from low to high CHIR, was dependent on Lef/Tcf and β-catenin transcriptional activity. These studies point to a non-transcriptional mechanism for β-catenin in regulation of NPCs, and potentially other stem progenitor cell types. Further, analysis of the β-catenin-directed transcriptional response provides new insight into induction of nephrogenesis.

**Summary Statement:** The study provides a mechanistic understanding of Wnt/ β-catenin activity in self-renewal and differentiation of mammalian nephron progenitors.

## Introduction

Wnt signaling is required for the self-renewal and differentiation of many key progenitor and stem cell type (Clevers et al., 2014). In canonical Wnt signaling, β-catenin Ctnnb1/β-catenin transforms Wnt ligand binding to cell surface receptor complexes into a transcriptional output (Clevers et al., 2014). Stabilization of β-catenin, in response to canonical Wnt signaling through Frizzled receptor complexes, results in the translocation of β-catenin to the nucleus, where β-catenin interaction with Lef/Tcf DNA-binding partners at cis-regulatory modules results in the transcriptional activation of target gene transcription (Cadigan and Waterman, 2012; Mazzotta et al., 2016). In a non-transcriptional role, β-catenin is critical in cadherin-mediated cell adhesion at adherence junctions. In this capacity, β-catenin is critical for the formation and morphogenesis of epithelia, cell-cell recognition, and the sorting of cell populations directed by distinct cell-surface cadherin complexes (Brembeck et al., 2006; Valenta et al., 2012).

The generation of a species appropriate number of nephrons – approximately 14,000 in the mouse and one million in the human kidney – is critically dependent on balancing the self-renewal and differentiation of nephron progenitor cells (NPCs; McMahon, 2016). In these events, Wnt9b-directed canonical Wnt signaling through β-catenin is implicated in both the control of the NPC self-renewal state (Brown et al., 2015; Carroll et al., 2005; Karner et al., 2011a; Park et al., 2007; Ramalingam et al., 2018a; Tanigawa et al., 2016) and the counter process of differentiation of NPCs (Karner et al., 2011b; Lindström et al., 2015; Park et al., 2007; Park et al., 2012; Ramalingam et al., 2018b; Tanigawa et al., 2016). In this inductive process, Wnt9b initiates a transcriptional program of active nephrogenesis and the associated cellular transition of mesenchymal NPCs into an epithelial renal vesicle, the precursor for each nephron (McMahon, 2016).

The development of an *in vitro* model for the culture of mouse NPCs enables NPC regulation to be studied in a controlled environment mirroring properties of the normal nephrogenic niche (Brown et al., 2015; Li et al., 2019; Tanigawa et al., 2015; Tanigawa et al., 2016). In nephron progenitor expansion medium (NPEM; Brown et al., 2015), Wnt signaling input is indirectly controlled by modulating β-catenin levels through varying levels of CHIR99021 (hereafter referred to as CHIR; Bennett et al., 2002), a small molecule inhibitor of glycogen synthase kinase-β (GSK3β): GSK3β directed phosphorylation of β-catenin controls cytoplasmic levels of β-catenin through the activity of a destruction complex (Zhang et al., 2023). *In vivo* and *in vitro* analysis of NPCs has demonstrated β-catenin associated at enhancers linked to genes active within NPCs or activated on activation of nephrogenesis (Karner et al., 2011; Park et al., 2012; Ramalingam et al., 2018; Guo et al., 2021). Genome scale chromatin interaction studies, and subsequent validation of predicted Lef/Tcf/ β-catenin-dependent cis-regulatory modules, provide strong support for β-catenin in transcriptional activation of target genes, in the induction of NPCs Park et al., 2012; Guo et al., 2021). In contrast, the data are less clear for β-catenin in NPC maintenance programs. While a focus on individual genes linked to the NPC state have provided support for a direct transcriptional role (Karner et al., 2011; Ramalingam et al., 2018) genomic scale analyzes have failed to provide a clear link to β-catenin dependent processes in NPCs, such as the control of cell proliferation (Guo et al., 2021).

To gain a deeper understanding of β-catenin’s diverse roles in NPC biology, we developed an approach for rapid, RNA-mediated genetic modification within NPCs *in vitro* and applied the strategy to studying the effects of modulating β-catenin and Tcf/Lef factor activity on the self-renewal and induction of primary mouse NPCs. Our findings point to a non-transcriptional role for β-catenin in the maintenance and expansion of NPCs and provide new insight into an early inductive pathway mediated through β-catenin and Lef/Tcf transcriptional complexes. These findings may have broader significance given the predominant role for Wnt regulation of stem/progenitor programs in metazoan (Lien and Fuchs, 2014; Loh et al., 2016; Rim et al., 2022).

## Results

### NPEM culture provides a rapid and consistent method for studying Wnt supported NPC self-renewal and differentiation

To model the self-renewal and differentiation of NPCs *in vitro*, primary mouse NPCs were isolated from embryonic day 16.5 (E16.5) kidneys by enzymatic digestion and magnetic depletion (**Fig. 1A**) and cultured in NPEM: NPEM is a defined medium formulated from insight into signaling pathways implicated in maintaining NPCs *in vivo* in the nephrogenic niche (Brown et al., 2015). NPC expansion over multiple generations requires low levels (1.25 µM) of the GSK3β inhibitor CHIR. As with NPCs *in vivo*, NPCs in low CHIR culture exhibited high levels of Six2 indicative of the self-renewal state (Kobayashi et al., 2008; Self et al., 2006)(**Fig. 1B, C**). In the kidney, *Six2* is downregulated in conjunction with the upregulation Lef1, a canonical Wnt target and transcriptional effector of Wnt signaling, and the Notch ligand Jag1, a critical regulator of early nephron patterning (**Fig. 1B**). In the developing kidney, Lef1+/Jag1+ cells initially aggregate, then undergo a mesenchymal-to-epithelial transition to establish an epithelial renal vesicle (RV), the anlagen for an adult nephron. Similarly, elevating CHIR levels (5 µM) in NPEM induced the differentiation of NPCs as evidenced molecularly by the up-regulation of Lef1 and Jag1and morphologically by the formation of tight cellular aggregates of induced NPCs (**Fig. 1C**).

**Figure 1:**
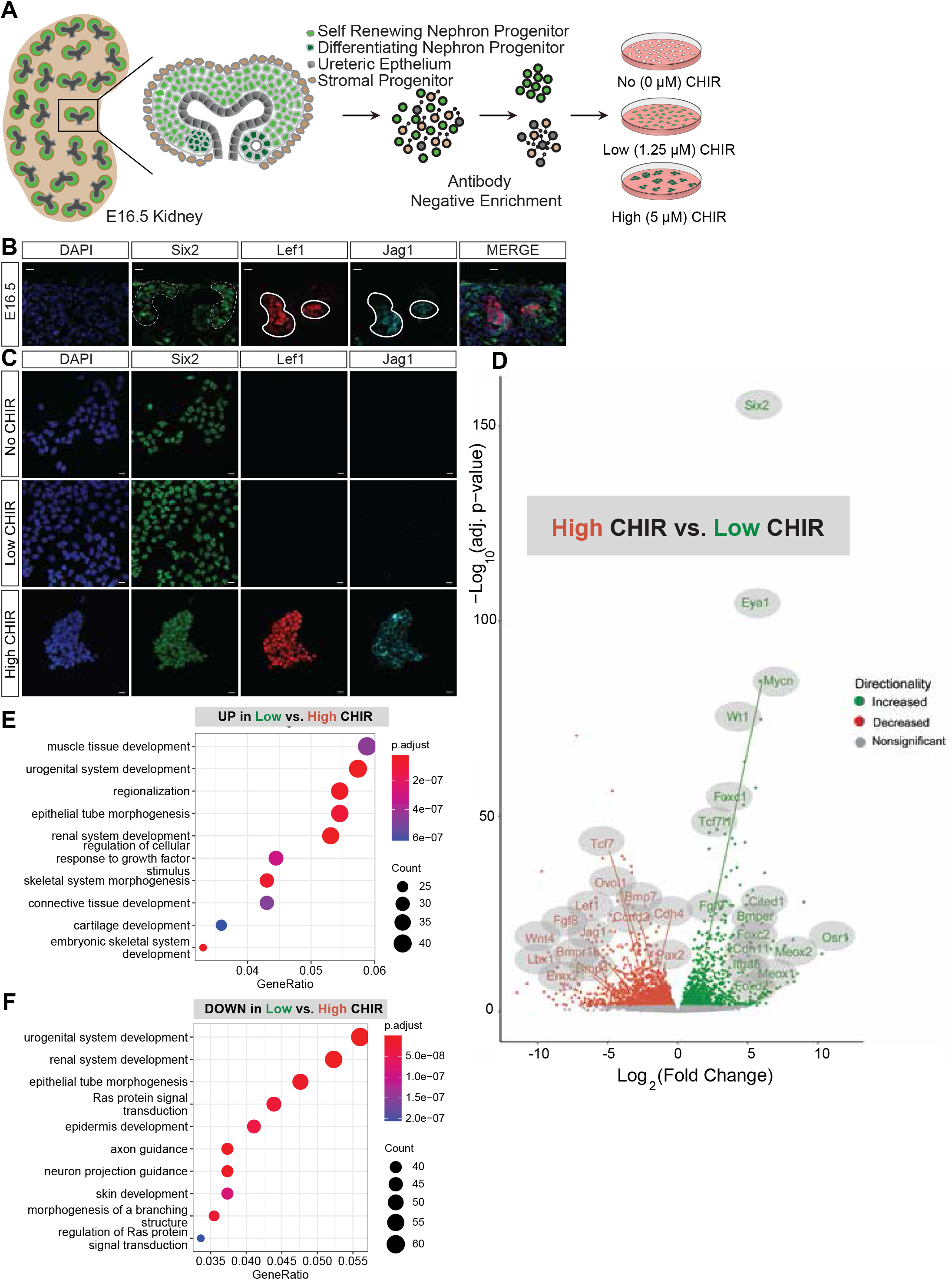
NPEM culture provides a rapid and consistent method for studying Wnt supported NPC self-renewal and differentiation. A) Schematic representation of NPC isolation and culture in NPEM supplemented with no (0μM) CHIR, low (1.25 μM) CHIR and high (5 μM) CHIR. B) Immunofluorescent staining for Six2, Lef1, Jag1, DAPI in E16.5 WT kidney (scale bar = 10 μm). Dotted lines highlight self-renewing area of the nephrogenic zone marked by high levels of Six2 protein and continuous line marks differentiating NPCs in the nephrogenic zone with high levels of Lef1 and Jag1. C) Immunofluorescence staining Six2, Lef1, Jag1, DAPI on WT nephron progenitor cells culture in no (0μm), low (1.25 μm) and high (5 μm) CHIR) (scale bar = 10 μm). D) Bulk RNA-seq. data of DEGs using Log2FC = absolute value cut off 0.5 and p-adjusted value cut off = 0.05 comparing low (1.25 μm) vs high (5 μm CHIR) post 24 hours of culture represented as a volcano plot. E) Top 10 most significant GO analysis of DEGs that are upregulated in low (1.25 μm) CHIR compared to high (5 μm) CHIR F) GO analysis of DEGs that are upregulated in high (5 μm) CHIR compared to low (1.25 μm) CHIR

To examine CHIR-mediated transcriptional responses, we performed a time course (0-12 h) and assayed key NPC self-renewal genes (*Six2, Cited1, Eya1*) and differentiation genes (*Wnt4, Lef1, Jag1, Fgf8* and *Lhx1*) by quantitative polymerase chain reaction (qPCR) to measure the transcriptional response to CHIR mediated Wnt activation (**Fig. S1A**). When CHIR was removed from NPC culture, we observed no loss of self-renewal-associated markers, rather expression was reproducibly elevated in the absence of CHIR (**Fig. S1A**) in agreement with earlier studies (Guo et al., 2021). In contrast, high CHIR had a marked and rapid (within 3hrs) effect on the transcriptional state of NPCs: a progressive downregulation in the expression of self-renewal cell markers and an elevation in the expression of genes linked to the induction of NPCs (**Fig. S1A**). The response was similar at low or high seeding density (37,500 to 300,000 cells/well; **Fig. S1B**) and correlated with an enhanced accumulation of β-catenin between tightly packed cell aggregates in high CHIR (**Fig. S2A**). To confirm responses reflected NPC actions, we evaluated the purity of NPC isolation by crossing an NPC specific CRE strain (Six2-GFP:CRE; Kobayashi et al., 2008) to tdTomato CRE reporter mouse (Madisen et al., 2010). Over 92% of cultured cells showed tdTomato, indicative of NPCs (**Fig. S1C**). Thus the measured responses reflected NPC gene activity.

To confirm low and high CHIR modelled responses reflected Wnt-pathway associated activity, NPCs were cultured with a bi-specific antibody (BSAB) that binds both Frizzled and LRP components of the Wnt-receptor complex, mimicking Wnt ligand signaling (Janda et al., 2017). NPCs cultured in 500 pM BSAB showed statistically significantly elevated cell proliferation (p value < 0.0001) as measured by EDU incorporation, but no evidence of an inductive response (**Fig. S1D, E**). In contrast, addition of 4 nM BSAB resulted in a marked down-regulation of the NPC marker *Cited1*, and elevated expression of *Jag1* and *Wnt4*, similar to high CHIR conditions (**Fig. S1D**), although the kinetics of *Six2* down-regulation differed from high CHIR (**Fig. S1D**). Collectively, these results indicate CHIR replicates key features of Wnt receptor action in NPC programs (**Fig. S1D, E**).

To examine CHIR-mediated responses in more detail, bulk RNA-seq profiling was performed on NPC cultures. NPCs were stabilized for 24 hours in low CHIR, then cultured for an additional 24 hours in the absence of CHIR, low CHIR or high CHIR conditions, before transcriptional profiling. Differential gene expression analysis using DE-SEQ2 (Log2 Fold-Change [Log2FC] cut off 1.0; adjusted p-value cut off 0.05) identified 824 DEGs up-regulated and 1205 DEGs down-regulated in the low to high CHIR transition (Table 1.1) and 603 DEGs up-regulated and 384 DEGs down-regulated on removal of CHIR (No CHIR) comparing with low CHIR conditions (Log2 Fold-Change [Log2FC] cut off 0.5; adjusted p-value cut off 0.05; Table 1.2). Comparing high CHIR to low CHIR showed a marked downregulation of genes encoding regulatory factors and genes specific to the NPC self-renewal state (**Fig. 1D; Table S1.1**), and the activation of genes associated with NPC differentiation (**Fig. 1D; Table S1.1**). Consistent with biological processes at play, Gene Ontology GO terms highlight nephron epithelium development and kidney epithelium development as significantly enriched terms for genes down-regulated in high CHIR (**Fig. 1E,F; Table S1.3**) and urogenital system development, renal system development, and epithelial tube morphogenesis amongst the top 10 most significant terms for the up-regulated gene set (**Fig. 1E,F; Table S1.4**).

In contrast, GO term analysis of low and NO CHIR highlighted altered cytoskeletal organization and attenuated cell movement in the absence of CHIR while the addition of CHIR results in a marked increase in cell proliferation associated terms in agreement with earlier studies (**Table S1.5-1.6**; Guo et al., 2021). Interestingly, in the absence of CHIR, transcript levels were actually elevated for several transcriptional determinants associated with the NPC self-renewal state (e.g. *Osr1, Meox1, Meox2*, and *Cited1*), consistent with earlier qPCR analysis **(Fig. S1A; Table S1.2**). Thus, Wnt/β-catenin signaling attenuates NPC gene activity arguing further against a transcriptional role for β-catenin in maintaining the NPC state. Elevated transcript levels may be a secondary consequence of reduced rates of cell proliferation in NPCs cultured without CHIR.

To determine whether the primary inductive response requires new protein synthesis, we switched NPCs from low to high CHIR in the presence of cycloheximide (CHX), an inhibitor of eukaryotic translation elongation (Burgers and Fürst, 2021; Schneider-Poetsch et al., 2010; Siegel and Sisler, 1963). Transcript levels of differentiation associated genes were up-regulated, albeit to lower levels than high CHIR alone, while the absence of *de novo* Jag1 indicated effective translational inhibition by CHX (**Fig. S2 B, C, D**). Interestingly, high CHIR and CHX treated NPCs failed to down-regulate key genes associated with the naïve NPC state such as *Six2, Cited1* and Eya1 consistent with a transcriptional and/or translational requirement for exit from the NPC program (**Fig. S2 B,C,D**). Retention of the NPC program may factor into the weaker inductive response observed in CHX.

### β-catenin promotes NPC proliferation but not the transcriptional program of self-renewing NPCs

To examine the role of Wnt-directed transcriptional mechanisms in NPCs, we developed and validated a novel platform for rapid, genetic modification of NPCs by mRNA lipofection (**Fig. 2A**). Lipofectamine (OPTI-mem) transfection of NPCs with polyadenylated mRNA encoding a GFP reporter showed a rapid onset of protein production: ∼ 30% of cells exhibited GFP fluorescence 3 h after transfection (**Fig. S3A**) and normal responses to varying levels of CHIR **(Fig. S3B**). When two mRNAs were introduced, encoding GFP and mCherry, up to 80% of cells were co-labeled 24 hours post-infection; thus, transfection competent cells take up multiple mRNAs (**Fig. S3C, D**). To initial characterize gene-knockdown, NPCs were harvested from mice constitutively expressing a Cas9-GFP allele (25), and transfected with a gRNA targeting GFP and a mRNA encoding mCherry. Fluorescence activated cell sorting (FACS) analysis identified a 75% reduction in GFP signal 24h post-transfection, predominantly localized to mCherry+ NPCs (**Fig. S3E**). Collectively, these experiments validate mRNA transfection as an approach to modifying endogenous gene activity within identifiable NPCs.

**Figure 2:**
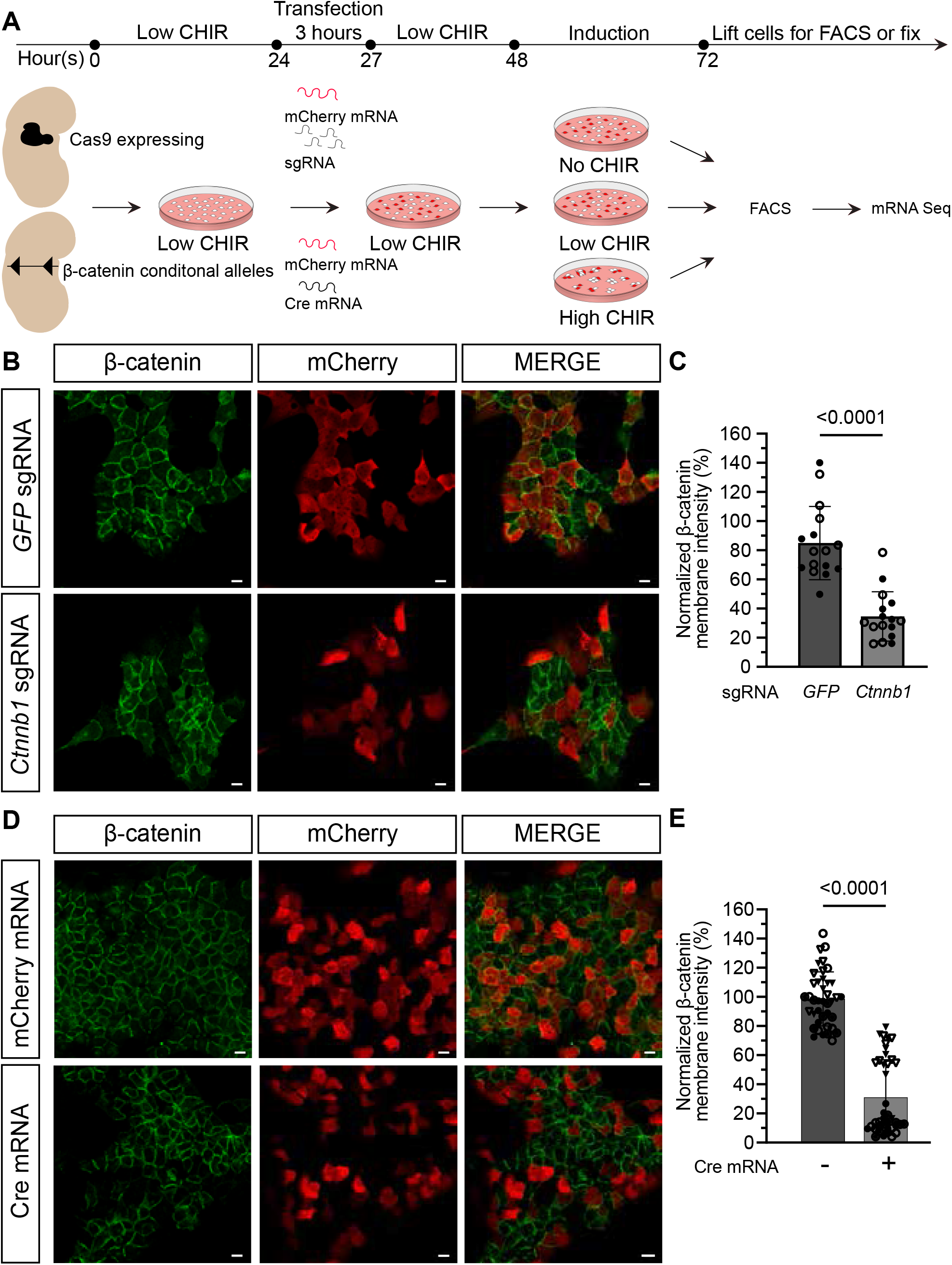
mRNA transfection provides rapid Cre or Cas9 mediated removal of β-catenin in primary mouse nephron progenitors. A) Schematic representation of Cre and Cas9 mediated transfection KO of NPCs from Cas9-GFP expressing and β-catenin conditional allele mice. NPCs are isolated, stabilized in 1.25 uM CHIR overnight (18-24 hours), transfected with either sgRNAs or Cre mRNA and mCherry mRNA and incubated in OPTI-MEM for 3 hours. Then NPCs are cultured in 1.25 μm CHIR for a 24-hour KO period prior to change in NPEM with differing CHIR levels. NPCs are assayed for bulk RNA-seq as well as immunostaining post 24 hours. B) Immunostaining of NPCs post 24-hour Cas9 mediated KO prior to media change into NPEM with differing levels of CHIR. β-catenin in green channel and mCherry (denoting transfected cells) in red (scale bar = 10 μm). C) Quantification of the reduction in membrane β-catenin protein signal post 24 hours Cas9 KO in transfected (mCherry+) NPCs. (n = 2 technical replicates, no biological replicates; 8 field of view/well; Mann-Whitney test). D) Immunostaining of NPCs post 24-hour Cre mediated KO prior to media change into NPEM with differing levels of CHIR. β-catenin in green and mCherry (denoting transfected cells) in red (scale bar = 10μm). E) Quantification of the reduction in membrane β-catenin protein signal post 24 hours Cre KO period in transfected (mCherry+) NPCs. (n = 2 biological replicates; 2-3 technical replicates, 8-10 fields of view/well; Mann-Whitney test).

To gain a deeper understanding into Wnt/β-catenin signaling, we utilized two different mRNA directed approaches to knock-out (KO) β-catenin activity (**Fig. 2A**): (1) KO of β-catenin (*Ctnnb1*) through targeted gRNA transfection into NPCs constitutively producing Cas9 and GFP (Platt et al., 2014), and (2) KO through Cre mRNA-mediated removal in NPCs homozygous for a conditional allele of β-catenin (Brault et al., 2001). In all experiments, co-introduction of an mCherry reporter mRNA was used to distinguish transfected cells and a guide RNA targeting GFP was used as a control in Cas9-directed KO studies.

Genetically appropriate NPCs were isolated from E16.5 kidneys, stabilized in low CHIR overnight and transfected for 3 h in OPTI-mem (**Fig. 2A**). To enable genetic removal and turnover of mRNA/proteins, NPCs were cultured in low CHIR for 24h, then cultured for an additional 24h in either low or high CHIR. Immunostaining showed β-catenin levels were reduced significantly in both gRNA- and CRE-directed KO conditions: ∼65% and ∼70%, respectively (**Fig. 2B-E**).

In low and high CHIR, removal of β-catenin resulted in a reduction in Edu incorporation resembling no CHIR conditions supporting the conclusion β-catenin promotes the proliferation of NPCs (**Fig. 3 A-C; Fig. S1E**) (Guo et al., 2021). Bulk RNA-seq was performed on reporter positive KO and control cells and the datasets intersected to identify a set of genes for individual conditions (**Table S1.7-Table S1.8**) and a robust set of DEGs shared between the two KO approaches (**Table S1.9, Table S1.10**). In low CHIR progenitor maintenance conditions, 13 genes were up regulated and 12 down-regulated on β-catenin removal; no significant alteration was observed in either the NPC transcriptional or proliferative transcriptional signatures (**Fig. 3D, E**; **Table S1.9-Table S1.10**). No kidney relevant GO terms associated with the downregulated gene-set (**Table S1.11**). However, GO term analysis of genes increased upon Cre and Cas9 β-catenin removal in low CHIR identified a list of 12 terms with kidney development (*Mmp9/Bcl2l11/Angpt2*) amongst the enriched terms reflecting an elevation of specific progenitor gene expression as observed on CHIR removal **(Table S1.12)**.

**Figure 3:**
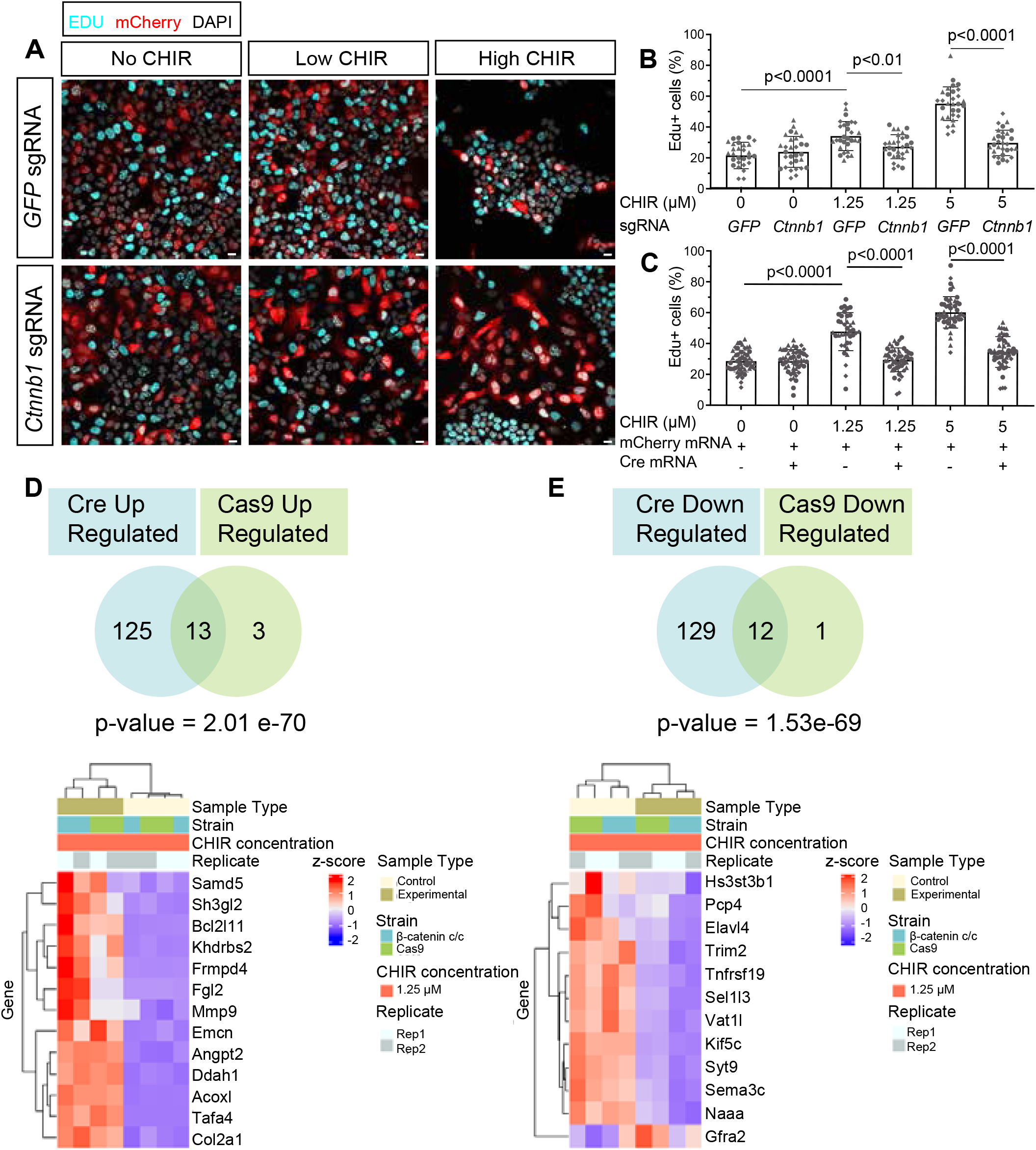
β-catenin promotes NPC proliferation but is not a transcriptional regulator of the self-renewal program. A) Edu chasing of NPCs with Cas9 mediated β-catenin removal post 24-hour culture (scale bar = 10 μm). B) Quantification of Edu+ (proliferating cells) per total number of NPCs in Cas9 β-catenin KO samples (9-10 fields of view/coverslip, 3 biological replicates all statistical tests are unpaired Student t test) C) Quantification of Edu+ (proliferating cells) per total number of NPCs in Cre β-catenin KO samples 3 biological replicates, 7-10 fields of view/coverslip, 5 technical replicates (except 1.25 CTRL, =4), from left to right: Mann-Whitney-test twice, unpaired t test, D) Intersection of DEGs upregulated (Cre up and Cas9 up) from NPCs with β-catenin removed in low CHIR. DEGs calculated with DESEQ2 using with p-adjusted cut offs = 0.05 and Log2FC cut off = 0.5. Heatmap of unbiased ranking based on significance of all upregulated genes compared to control low CHIR samples of the intersection of both Cre and Cas9 mediated removal of β-catenin in low CHIR E) Intersection of DEGs down regulated (Cre down and Cas9 down) from NPCs with β-catenin removed in low CHIR. DEGs calculated with DESEQ2 using with p-adjusted cut offs = 0.05 and Log2FC cut off = 0.5. Heatmap of unbiased ranking based on significance of all downregulated genes compared to control low CHIR samples of the intersection of both Cre and Cas9 mediated removal of β-catenin in low CHIR

Intersection of the gene set exhibiting a shared down regulation in low CHIR with β-catenin/Tcf7/Lef1 ChIP-seq data specific to low CHIR from published data (283 associations ;Guo et al., 2021) showed no overlap between the datasets. Earlier reports have suggested a direct Wnt/β-catenin regulation of a number of NPC self-renewal associated genes including Tafa5 (Karner et al., 2011b). However, expression of these genes was not altered on β-catenin removal in low CHIR. Examining ChIP-seq analysis of β-catenin/Tcf/Lef interactions in primary NPCs in published datasets (Guo et al., 2021) demonstrated Tafa5 undergoes a similar Tcf/Lef binding switch as NPC differentiation genes *Wnt4* and *Jag1*suggesting Tafa5 is potentially a direct target of β-catenin-mediated transcriptional regulation in differentiating NPCs (**Fig. S4A**). Collectively, these data lend additional support to the view that β-catenin does not play a direct transcriptional role in either NPC maintenance or proliferative programs.

### β-catenin removal maintains the nephron progenitor state in the presence of inductive signaling

Under high CHIR conditions, β-catenin KO resulted in the persistence of Six2 and the repressive Lef/Tcf regulatory factor Tcf7l1 which is normally down-regulated on induction of NPCs (Guo et al., 2021), a failure of Jag1 induction and the absence of cell aggregation (**Fig. 4A-E; Fig. S5 A-F**). In short, β-catenin KO NPCs in high CHIR resembled NPCs maintained under low CHIR conditions. Consistent with this view, principal component analysis (PCA) showed co-clustering along the PC1 axis for NPCs in all low CHIR conditions with NPCs having undergone β-catenin removal and cultured in high CHIR conditions (**Fig. S6A**). Examining DEGs comparing both β-catenin removal conditions in high CHIR identified a highly significant response (hypergeometric test: p-value for up-regulated, 3.07e-1664; p-value down-regulated, 3.675e-776) with 937 shared DEGs: 282 down-regulated following β-catenin KO and 655 up-regulated (**Fig. 5A-B; Table S1.14, Table S1.15, Table S1.16, Table S1.17**). Ranked intersections of shared genes up regulated on KO in high CHIR identified *Cited1* as the most significant gene (p-value = 3.57E-36 ; Log2FC = 5.01) together with other well-known identifiers of an uninduced self-renewal NPC state including *Tmem100, Meox2* and *Osr1* in the top 30 ranked gene set (**Fig. 5C; Table S1.17**). NPC self-renewal genes comprised much of the highly ranked GO term “urogenital system development” recovered from GO analysis of the entire gene set (**Fig. 5E; Table S1.18**). Other terms such as “**ameboidal cell movement**”, “**epithelium migration**” and “**tissue migration**” likely reflect altered cell behaviors observed on induction of NPCs following β-catenin removal (**Fig. 5E; Table S1.18**).

**Figure 4:**
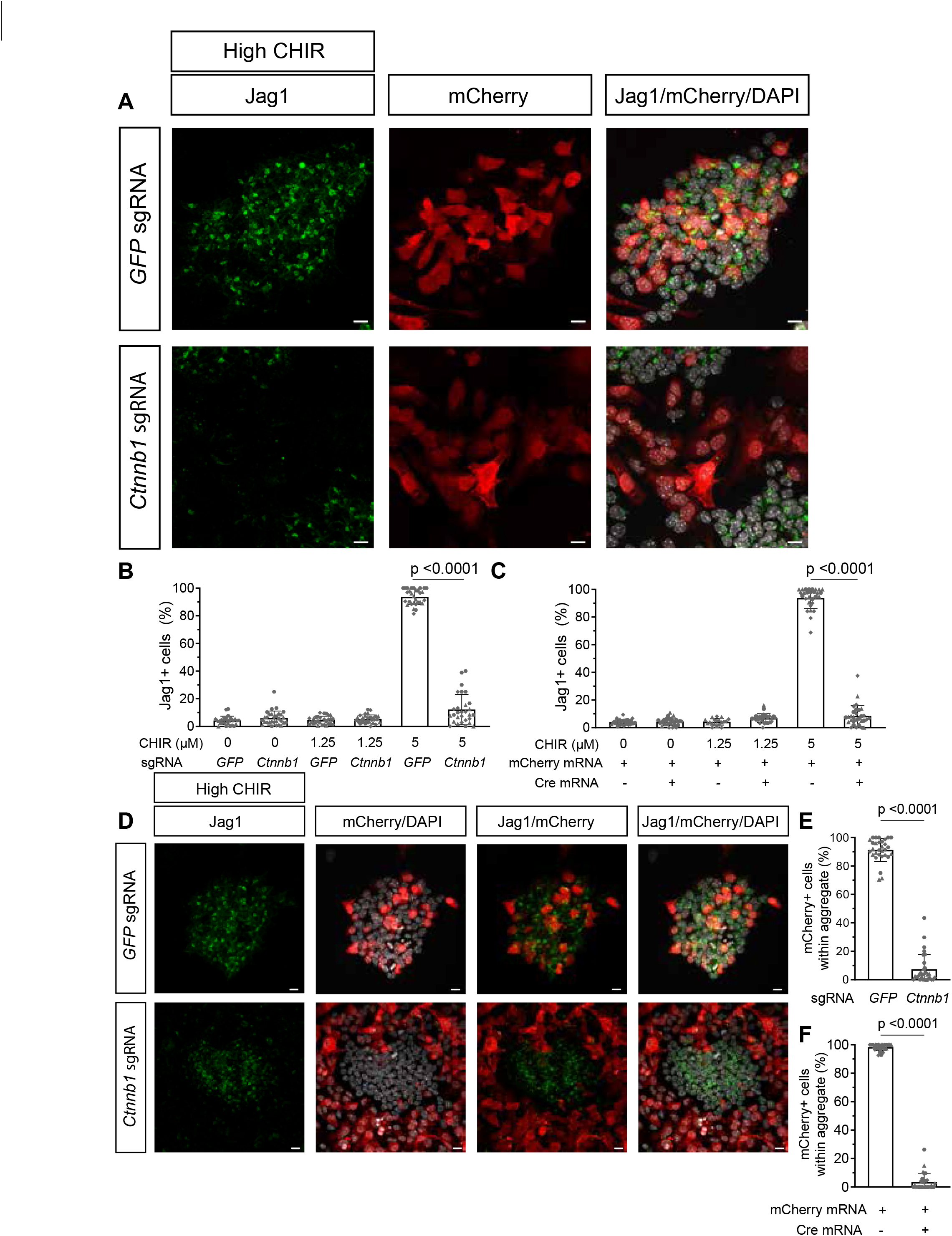
Induction status and cell behavior changes /resulting from β-catenin removal in nephron progenitors in high CHIR. A) Jag1 protein is lost in Cas9 mediated removal of β-catenin in NPCs in mCherry+ transfected cells in 5 μm CHIR. DAPI marks nuclei. B) Quantification of Jag1 protein in mCherry + NPCs in Cas9 mediated removal of β-catenin. 7-10 fields of view/coverslip, 3 biological replicates, normality test did not pass, Mann-Whitney test. C) Quantification of Jag1 protein in mCherry + NPCs in Cre mediated removal of β-catenin. 9-10 fields of view/coverslip, 2 biological replicates, normality test did not pass, Mann-Whitney test. D) NPCs with β-catenin removed (transfected cells) do not form tight aggregates. mCherry marks transfected cells, Jag1 induction marker and DAPI (nuclei marker). E) Percentage of mCherry+ transfected NPCs in NPCs with β-catenin removed (Cas9 method) within aggregates. 10 fields of view/3 biological replicates. normality test did not pass, Mann-Whitney test. F) Percentage of mCherry+ transfected NPCs in NPCs with β-catenin removed (Cre method) within aggregates. 9-10 fields of view/2 biological replicates, normality test did not pass, Mann-Whitney test.

**Figure 5:**
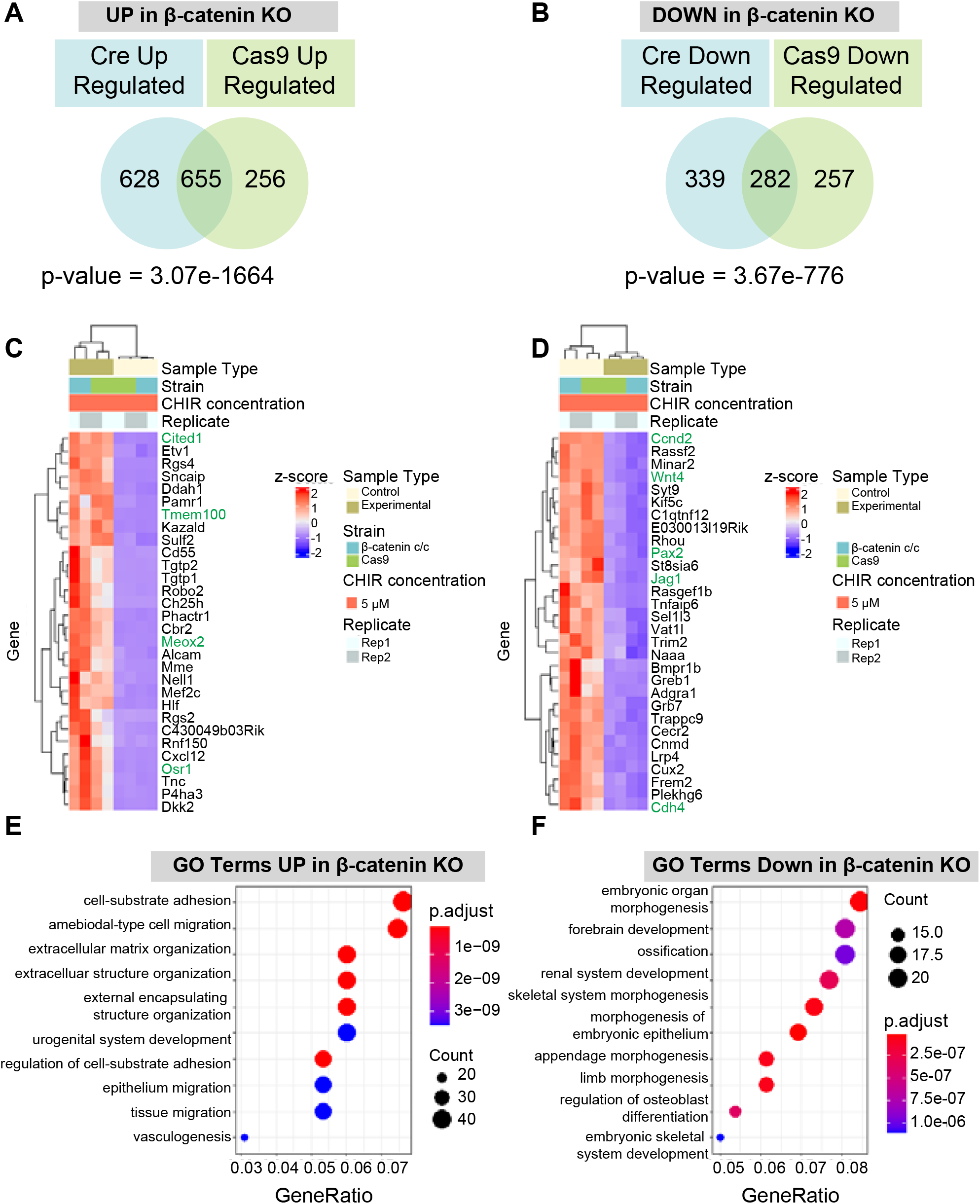
Transcriptional changes resulting from β-catenin removal in nephron progenitors. A) Intersection of DEGs upregulated from Cre and Cas9 mediated β-catenin removal B) Intersection of DEGs downregulated from Cre and Cas9 mediated β-catenin removal C–D) Heatmap of unbiased ranking of top 30 significant upregulated (C) and downregulated (D) genes compared to CTRL samples of the intersection of both Cre and Cas9 mediated removal of β-catenin in high CHIR. Green genes are known NPC self-renewal markers (C) or known NPC differentiation markers or canonical Wnt/β-catenin targets (D). Genes in green are previously identified markers of either self-renewal (C) or differentiation/ Canonical Wnt target (D). E) Top 10 most significant GO associated with genes upregulated as a result of β-catenin removal in high CHIR of intersected Cre and Cas9 meditated DEGs F) Top 10 most significant GO associated with genes downregulated as a result of β-catenin removal in high CHIR of intersected Cre and Cas9 meditated DEGs

### β-catenin transcriptional targets in the induction of nephron progenitors

Amongst the top 30 most significant down-regulated genes on KO of β-catenin in high CHIR were well characterized genes associated with the differentiation of NPCs including Wnt4 and Jag1 (**Fig. 5D**). and the cell cycle regulator, *Ccnd2*, a known target of Wnt-regulated cell proliferation (**Fig. 5D; Table S1.16**). As expected, top GO terms in this group of genes were associated with kidney developmental programs: “**embryonic organ morphogenesis**”, “**renal system development**” and “**morphogenesis of an embryonic epithelium**”; and Wnt signaling, including “**Wnt signaling pathway**” and “**cell-cell signaling by Wnt**”, consistent with a modified Wnt-signaling response (**Fig. 5E, F; Table S1.19**).

Tcf/Lef factors are direct transcriptional effectors binding regulatory DNA around Wnt target genes (Cadigan and Waterman, 2012). Of the four Tcf/Lef factors, fluorescent *in situ* hybridization and mouse scRNA-seq data analysis (Guo et al., 2021; Kim et al., 2023 preprint) point to elevated expression of *Tcf7l1*in self-renewing NPCs and *Lef1* following NPC induction. Tcf7 and Tcf7l2 expression did not change markedly between NPC states (**Fig. S7 A-C**). To assay the requirement for Tcf/Lef factors in NPC programs, we optimized individual gRNAs for the removal of each factor, extending the culture protocol for an additional 24h following sgRNA introduction (48h total) to ensure a strong knock-down of Tcf/Lef factors (**Fig. 6A**). Importantly, 24h of additional NPC culture did not alter gene expression trends for *Six2, Cited1, Wnt4* or *Jag1*, key anchor genes employed throughout the study (**Table S1.20**). Individual sgRNAs led to a removal of greater than 80% of Tc7l1, Tc7l2 and Lef1 and 50% of Tcf7 (**Fig. S8A-H**). When all four sgRNAs were combined in high CHIR conditions, we observed a ∼50% reduction of Jag1+ cells in Lef/Tcf KO cells quantifying Golgi-localized Jag1 to a general Golgi marker (**Fig. 6B, C**). Thus, KO of Lef/Tcf factors attenuated the CHIR-directed differentiation response consistent with β-catenin acting at least in part through a canonical Tcf/Lef transcriptional pathway.

**Figure 6:**
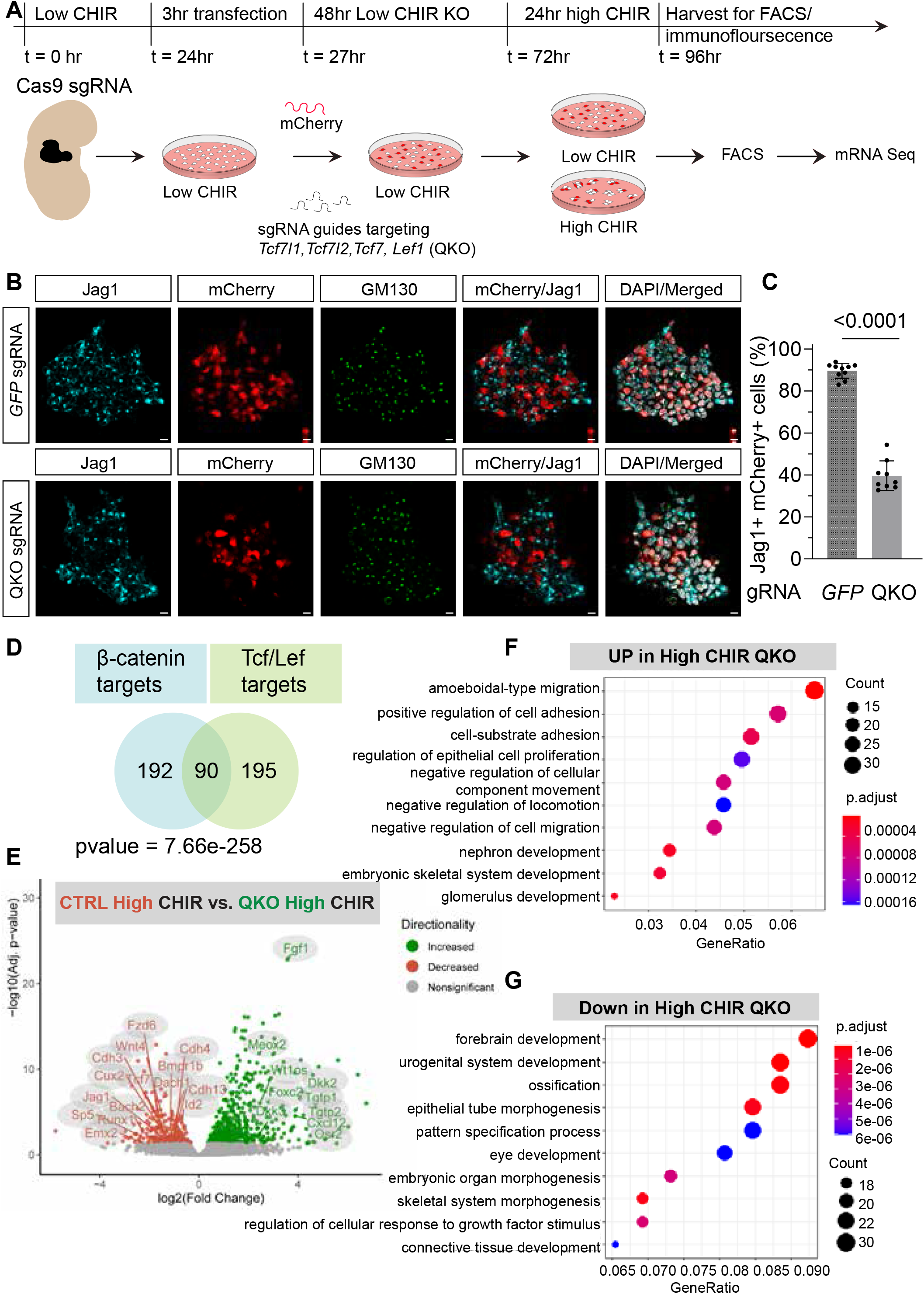
Tcf/Lef transcription factor removal in nephron progenitors enables β-catenin target validation. A) Schematic representation of Cas9 mediated removal of Tcf7l1, Tcf7l2,Tcf7 and Lef1 in NPCs isolated from Cas9-GFP expressing mice. NPCs were isolated, stabilized in 1.25 μm CHIR overnight (18-24 hours), transfected with sgRNAs (targeting Tcf7l1, Tcf7l2,Tcf7 and Lef1 in experimental conditions or GFP sgRNA in CTRL conditions) along with mCherry mRNA and incubated in OPTI-MEM for 3 hours. Then NPCs are cultured in 1.25 μm CHIR for a 48-hour KO period prior to change in NPEM with differing CHIR levels. NPCs are assayed for mRN-seq as well as immunostaining 24 hours later. B) Immunofluorescence staining of NPCs with Tcf7l1,Tcf7l2,Tcf7and Lef1 (QKO). Induction marker Jag1 = cyan, mCherry = red, Golgi = green marked by GM130 antibody, DAPI= gray. 10μm scale bar. C) Quantification of induction marker Jag1 in QKO NPCs cultured in 5μm CHIR. Unpaired T-test. 1 biological replicate, 8 fields of view. D) Intersection of QKO target genes (genes lost in QKO with Log2FC cut off 0.5 and p-adj cut offs 0.05 with Ctnnb1 KO target genes (genes lost in Ctnnb1 Cre and Cas9 KO with Log2FC cut off 0.5 and padj cut offs 0.05) E) Volcano plot of DEGs comparing QKO samples vs. control in high CHIR conditions. F) Top 10 most significant GO of DEGs upregulated in QKO samples in high CHIR conditions G) Top 10 most significant GO of DEGs downregulated in QKO samples in high CHIR conditions

PCA analysis of bulk mRNA-sequencing on genetically modified NPCs from the quadruple (*Tcf7l1, Tcf7l2, Tcf7, Lef1*) KO population (QKO) in high CHIR demonstrated QKO cells clustered closer to NPCs maintained in low CHIR conditions (**Fig. S7D**). As expected from β-catenin experiments, DEG analysis identified few differences between wild type (WT) and QKO (n=24) samples in low CHIR conditions, with all but one of the genes up-regulated on *Lef/Tcf* removal (**Table S1.21**). In contrast, a large number of DEGS (n=857) distinguished control and QKO NPCs in high CHIR (**Fig. 6E**). Of these, 572 were up-regulated and 285 down-regulated on *Lef/Tcf* removal **(Table S1.22)**. For this latter set requiring Lef/Tcf factors for the normal induction of gene activity, ∼30% were shared with the β-catenin removal set in high CHIR conditions (**Fig. 6D; Table S1.23)**, and as expected, shared similar GO terms with this gene set **(Fig. 6 E,F; Table S1.24-1.25)**. These data support the conclusion that β-catenin acts through Lef/Tcf factors in regulating the transcriptional inductive response initiating differentiation of NPCs.

To predict genes that were most likely direct Wnt/β-catenin targets within the NPC inductive response, we intersected the 282 β-catenin dependent genes from the dual genetic KO gene list with published ChIP-seq analysis of the chromatin association of β-catenin, Tcf7and Lef1 in high CHIR conditions (n =5011 associations **(Table S1.13**; Guo et al., 2021). A significant overlap (p-value = 2.94e-40) was observed with 161 genes using HOMER (Duttke et al., 2019) to assign genes to peaks within 100 kb of the transcriptional start site **(Table S1.26).** Intersecting these putative target genes with annotated single cell datasets of the developing (p0) mouse kidney undergoing early patterning from NPCs identified 137 of the 162 genes within developing mouse nephron *in vivo* (Kim et al., 2023) (**Table S1.27**). Examining this set in parallel data for the developing human kidney (Lindström et al., 2021) showed the majority (117) shared within similar populations in the developing human kidney (**Fig. 7A, Table S1.28**). These analyzes provide strong evidence that *in vitro* inductive responses in CHIR treated NPCs reflect *in vivo* activity of β-catenin directed, Lef/Tcf driven, transcriptional programs in committed NPC cell types.

**Figure 7:**
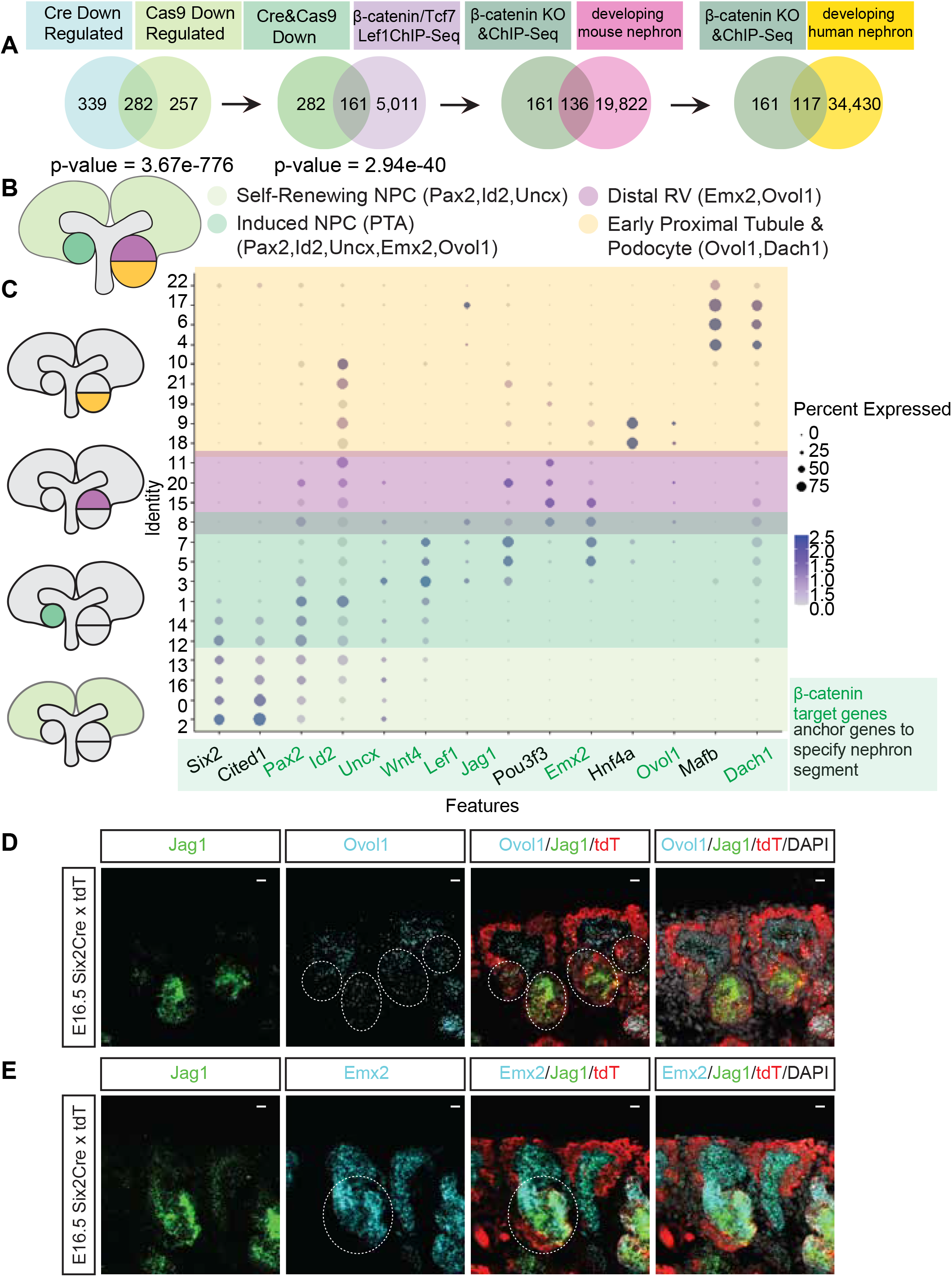
Integration of published ChIP-seq data and scRNA-seq data with β-catenin target genes reveals a suite of genes responsible for the early induction of NPCs and patterning of the developing nephron. A) Intersection of β-catenin target genes with β-catenin/ Tcf7/Lef1 ChIP-seq binding and integration with scRNA-seq data narrows down a significant list of 137 genes expressed in the developing nephron B) Schematic representation of the nephrogenic niche. PTA: pre-tubular aggregate, RV: renal vesicle. C) Dot plot denoting co-expression of β-catenin target genes Pax2, Id2, Uncx, Emx2, Ovol1, Dach1 with known markers of self-renewing NPCs (Six2, Cited1), early induction gene (Wnt4), distal marker (Pou3f3), proximal tubule marker (Hnf4a), podocyte gene (Mafb) in mouse p0 scRNA-seq D) RNA-scope of Ovol1 on E16.5 Six2TGCxTdt kidneys (10μm scale bar). Dotted lines denote Ovol1 transcripts in differentiating NPCs. E) RNA-scope of Emx2 on E16.5 Six2TGCxTdt kidneys. (10μm scale bar). Dotted lines denote Emx2 transcripts in differentiating NPCs.

β-catenin target genes linked to chromatin association with β-catenin and Tcf/Lef factor in high CHIR conditions include Ovol1 and Emx2 which encode transcriptional regulators that may play a role downstream of induction, in nephron patterning as Lef/Tcf factors are bound at putative cis regulatory elements at Ovol1 and Emx2 in NPCs cultured in High CHIR (Guo et al., 2021) (**Fig. S9 D, E**)). Fluorescent in situ hybridization demonstrated *Ovol1* and *Emx2* were expressed early in differentiating NPCs, overlapping with previously established NPC differentiation markers genes (**Fig. 7C-E**). Subsequently, *Ovol1*was expressed broadly within the developing renal vesicle while *Emx2* refined to distal regions (**Fig. 7C-E**).

## Discussion

Our analysis of β-catenin action in the self-renewal programs that lead to NPC expansion and the inductive program that initiates the process of nephrogenesis leads to several broad conclusions. (1) β-catenin does not transcriptionally regulate the NPC self-renewal state. However, low CHIR and β-catenin are required to support normal levels of NPC proliferation. (2) β-catenin orchestrates the differentiating response with Lef/Tcf binding factors indicating canonical Wnt transcriptional complexes play a key in the induction of NPCs. (3) Removal of either β-catenin or Lef/Tcf factors modifies cell behaviors with KO cells excluded from high CHIR-induced tight cell aggregates. Additional insight and support for this conclusion is presented in the accompanying paper (Der et al., 2023).

### The role of β-catenin in self-renewal of nephron progenitors

A number of genes have been reported to display Wnt9b-dependent expression in self-renewing NPCs (Karner et al., 2011b) linking Wnt9b to the maintenance of NPC programs and properties (Carroll et al., 2005). Though seven of the 16 reported Wnt9b-regulated genes in these studies showed elevated expression of at least one isoform in low CHIR versus no CHIR analyzes in previous reports (Guo et al., 2021), none of these DEGs showed differential expression upon β-catenin removal in our data. One of these, *Tafa5/Fam19a5*, has been shown to exhibit Lef/Tcf binding at putative regulatory regions (Guo et al., 2021; Karner et al., 2011b). Thus, *Tafa5* may be an early target of the inductive response.

Direct analysis of NPC cultures in the absence of CHIR, genetic removal of β-catenin and *Lef/Tcf* activity, and Tcf7/Lef/β-catenin chromatin association studies (Guo et al., 2021) argue against a role for the canonical Wnt/β-catenin transcriptional pathway in control of the naïve NPC state, preceding Wnt-mediated induction. Additional support for this conclusion comes from time-lapse imaging of kidney explants from a Lef/Tcf-GFP reporter mouse, where canonical pathway activity was restricted to differentiating NPCs (Lindström et al., 2015). β-catenin is known to associate with other transcriptional partners outside of canonical Lef/Tcf transcriptional regulators and activate transcription (Doumpas et al., 2019; Mukherjee et al., 2022). However, β-catenin removal is still expected to identify genes directly linked to regulation of the NPC state in low CHIR. Our studies all consider a single time point in the analysis of outcomes. Recent studies in the intestine point to different chromatin interactions at different time points after pathway activation (Borrelli et al., 2021). Though regulation of proliferation might be expected to be a broad program, and heterogenously regulated in a non-synchronized NPC population, a single time point analysis is a caveat to the conclusion of a non-transcriptional role of β-catenin in proliferative expansion of NPCs.

β -catenin is reported to play an essential, non-transcriptional role in self-renewal of mouse epiblast stem cells (Kim et al., 2013). The reduction of NPC proliferation, as measured by EDU pulse chase experiments on removal of CHIR or KO of β-catenin in low CHIR maintenance conditions, points to an action for β-catenin in the proliferative expansion of NPCs in NPEM culture. How is not clear. Interestingly, β-catenin KO in low CHIR results in elevated levels of transcripts encoding key genes associated with the NPC progenitor state, including *Cited1*, *Six2*, *Meox1* and *Eya1*, and the repressive Wnt transcriptional regulatory factor Tcf7l1. These findings suggest β-catenin activity may actually attenuate NPC self-renewal through some unknown mechanism. Alternatively, as β-catenin is required for normal rates of NPC proliferation, reduced rates of cell division following β-catenin removal, either genetically or the removal of CHIR, may enhance accumulation of NPC transcripts and their protein products, given a lengthened period for transcription between each cell cycle. Further clarification will benefit from additional study of β-catenin in NPCs focused on non-transcriptional mechanisms.

### β-catenin/Tcf/Lef transcriptional targets

Examining the inductive response *in vitro* provides new insight into Wnt/ β-catenin targets in the early activation of the nephrogenic program. Many of these intersect with independent studies predicting transcriptional targets through chromatin association studies examining β-catenin and Lef/Tcf factors in NPCs *in vivo* (Park et al., 2012) and *in vitro* in NPEM culture (Guo et al., 2021). Lef/Tcf removal studies, in conjunction with β-catenin KO, indicate that β-catenin acting in partnership with Lef/Tcf factors – canonical Wnt transcriptional activity - is a major mechanism underpinning β-catenin-mediated induction of NPCs.

Our approach to analysis of NPC induction pathway *in vitro* enables a tight control on the experimental conditions, identifying known targets, and predicting new targets through intersection of chromatin binding and KO studies. The transcription factors Cux2, Ovol1, Sox11, Bach2, Id2 and Emx2 are predicted to be directly regulated through /Tcf/Lef/ β-catenin interactions consistent with studies in other systems (Hasenpusch-Theil et al., 2018; Kormish et al., 2010; Mahmoudi et al., 2009; Piloto and Schilling, 2010; Rockman et al., 2001; Schmidt-Ott et al., 2007). Interestingly, *Emx2* and *Ovol1* are both expressed in the distal regions of the developing nephron, in both the developing mouse and human nephron anlagen. *Ovol1* and *Emx2* mutant mice have reported phenotypes of embryonic cystic kidneys and urogenital defects, respectively (Miyamoto et al., 1997; Teng et al., 2007). However, we were unable to observe cystic, or other kidney phenotypes, in *Ovol1* mutant mice generated through gRNA-directed KO studies (HB and APM, unpublished studies).

We observed a β-catenin dependent up-regulation of *Bmp7* and *Bmpr1b* in high CHIR conditions consistent with Wnt-regulation of Bmp signaling in the nephrogenic program. Several lines of evidence support Wnt and Bmp interactions in the induction of NPCs. BMPs have been shown to play an important role in NPC self-renewal and differentiation (Brown et al., 2013; O’Brien, 2019; Oxburgh et al., 2011; Oxburgh et al., 2014). Inductive Wnt/β-catenin signaling is proposed to act upstream of Bmp signaling, and *Bmp7* is expressed in RV’s overlapping *Jag1* and *Wnt4* (Park et al., 2012). *Wnt4* is itself an inductive signal, maintaining the initial Wnt9b-triggered response as an autoregulatory signal (Stark et al. 1994), transcriptional regulated by canonical Wnt-regulatory complexes (Park et al., 2012). *Bmp7* enhancers are bound by β-catenin at predicted Lef/Tcf binding motifs. Further, *Bmp7* enhancer activity is lost on mutation of Lef/Tcf binding sites within cis-regulatory modules regulating Bmp7 activity in developing nephrons (Park et al., 2012). Bmp7 has been proposed to signal through Bmpr1b from analysis of scRNA-seq data from the developing mouse kidney (Combes et al., 2019). Wnt targeting of signaling pathways may enable cell interactions critical for epithelialization and patterning of the early nephron precursor.

In addition to *de novo* induction of genes in high CHIR, β-catenin activity was required to silence expression of genes associated with the NPC self-renewal state, including *Six2* and *Cited1*. Interestingly, cyclohexamide studies suggest that while the early inductive program was activated in high CHIR, there was no reduction in the expression of self-renewal target genes. Thus, new protein synthesis is not essential for inducing key Wnt targets but is essential for silencing the NPC-associated regulatory program. Examining the list of putative β-catenin targets identifies the known transcriptional repressor *Id2*. *Id2* induction in high CHIR is perturbed on loss of β-catenin or QKO of Lef/Tcf factors. Id2 is a mediator of the BMP pathway suggesting a potential link between Wnt and Bmp signaling (Blank et al., 2009; Brown et al., 2013) and is reported to act alongside *Six2* in gut development (Mori et al., 2018). Mechanistically, Wnt/β-catenin targeted activation of an NPC inhibitory factor(s) could provide a rapid feed-forward activity destabilizing the progenitor state

### Canonical Wnt regulation in other stem/progenitor systems

Several well characterized developmental processes and regeneration in adult stem cell systems rely on canonical Wnt activity in regulation and organization of stem cell populations (Briscoe and Small, 2015). Hair follicle stem cells (HFSC) in the skin and intestinal stem cells (ISCs) in the gut (Clevers et al., 2014; Gehart and Clevers, 2019a; Lien and Fuchs, 2014) are some of the best studied examples. The HFSC system is transcriptionally similar to the NPCs. Tcf7l1 and Tcf7l2 are present in the quiescent Wnt (inactive or low) state. Upon canonical Wnt signaling input, “activating” Tcf7 and Lef1 factors invokes the transition of HFSCs to a highly proliferative state with activation of canonical Wnt targets (Guo et al., 2021; Lien and Fuchs, 2014; Lien et al., 2014; Merrill et al., 2001). Targeted removal of β-catenin from the HFSC population blocks Wnt-mediated induction of HFSCs (Choi et al., 2013; Lien et al., 2014), much as β-catenin removal from NPCs blocks CHIR-mediated induction of the nephrogenic program in vitro and Wnt-directed differentiation in the in vivo kidney (Park et al., 2007). Conversely, overexpression of β-catenin enhanced ectopic NPC differentiation in the kidney and hair follicle induction in the skin (Gat et al., 1998; Lowry et al., 2005; Närhi et al., 2008; Park et al., 2007).

In the mammalian gut, the Wnt pathway has been linked to stem cell maintenance, differentiation, and maintenance of the intestinal epithelium (Barker et al., 2007; Gehart and Clevers, 2019b; Sato et al., 2011; Yan et al., 2017). Loss of *Ctnnb1* and *Tcf7* functions in the gut leads to a dramatic loss of ISCs, reduction of cell proliferation and a disruption of epithelial organization and the differentiation of mature cell types (Korinek et al., 1998; Fevr et al., 2007). However in contrast to the NPC populations where a transcriptional Wnt/ β-catenin/Tcf activity is only responsible for the differentiation, a direct transcriptional Wnt/ β-catenin/Tcf mediated response regulates the self-renewal and differentiation of ISCs (Fevr et al., 2007; Guo et al., 2021; Korinek et al., 1998). In this, β-catenin interactions with Tcf7l2 transcriptionally regulates the self-renewal and differentiation of healthy and malignant intestinal stem cells (van de Wetering et al., 2002). Moreover, fine tuning of a Wnt mediated response in the self-renewal and differentiation of ISCs has been recently attributed to the differential binding of β-catenin to either N-terminal or C-terminal co-factors in the nucleus directing distinct transcriptional outcomes (Borrelli et al., 2021).

In summary, the current study highlights the complementary insight that can be obtained into complex biological processes in the developing kidney from a well-controlled *in vitro* system that replicates key features of renal development. The current studies raise several important questions merit further study: the non-transcriptional role of β-catenin in NPC proliferation; inductive programming of morphological and patterning responses; and, the potential for transcriptional feedback into silencing of the NPC program. Further, the *in vitro* approach and NPC systems is a powerful platform for deepening a mechanistic understanding of the regulatory processes balancing maintenance, expansion and commitment of stem/ progenitor cells, and the mechanistic action of Wnt, and other, regulatory inputs (Gehart and Clevers, 2019a; McMahon, 2016; van Es et al., 2012).

## Materials and Methods

See Table S2 for mRNA synthesis primers, CRISPR sgRNA sequences, RT-qPCR primers, RNA-Scope probes, small molecules and utilized antibodies. Bulk mRNA sequencing Data will be available on GEO. GSE232482 for mouse p0 scRNA-seq.

### NPC isolation and culture

NPEM formulation and NPC isolation followed the published protocol (Brown et al., 2015). Kidneys were harvested from E16.5 mice embryos and placed into PBS on ice. Each kidney was expected to yield approximately 150,000 NPCs (100,000 NPCs for B6 background). After dissection, kidneys were washed with 2 ml HBSS (Thermo Fisher Scientific, 14175-095) twice to remove blood and shaken on a Nutator platform for 2 min at 495 rpm, then incubated in 2 ml HBSS solution containing 2.5 mg/ml collagenase A (Roche, 11 088 793 001) and 10 mg/mL pancreatin (Sigma, P1625) for 15 min at 37℃ while rocking on a Nutator platform at 495 rpm. The nzymeatic reaction was then terminated by the addition of 125 μl of fetal bovine serum (FBS). The resulting supernatant was pelleted then passed through a 40 μm filter, and then washed with AutoMACS running buffer (Miltenyi, 130-091-221) before spinning down at 500g for 3 min. The cell pellet, predominantly cells of the cortical nephrogenic zone, was resuspended in 76 μl of AutoMACS running buffer from 10 pairs of kidneys. NPC enrichment resulted from the removal of other cell types in the cell suspension using a combination of PE-conjugated antibodies as follows:

- Anti-CD105-PE (Miltenyi, 130-102-548), 9 μl
- Anti-CD140-PE (Miltenyi, 130-102-502), 9 μl
- Anti-Ter119-PE (Miltenyi, 130-102-893), 8 μl
- Anti-CD326-PE (Miltenyi, 130-118-075), 1.6 μl

The cells and antibodies were incubated at 4 ℃ for 30 min without agitation on ice, then washed three times with 1 mL AutoMACS running buffer. To remove the unwanted non NPCs, 20 μl of anti-PE beads were added to the cell suspension for 30 min on ice. Cells were then washed three times in 1 ml of running buffer, and finally cells resuspended in 1 ml of AutoMACS running buffer and sent through the AutoMACS program as described in the published protocol to remove non-NPC cell types enriching for NPCs..

NPCs were seeded at 300k cells/well on a 24 well plate. 24-well NPC culture plates were treated with Matrigel (Corning, 354277) 1:25 in APEL medium and incubated at room temperature in cell culture hood for at least 1 hr. APEL was then aspirated off leaving behind remaining Matrigel. For all culture experiments, NPCs were seeded in low (1.25μM) CHIR NPEM. Upon seeding, plates were shaken three times every 10 minutes to evenly distribute cells throughout the well.

### Cycloheximide (CHX) experiments

CXH (14126, Cayman Chemical) was resuspended in DMSO and stored as 25 mg/ml stock solution at - 20 C. To achieve the final 10μM working concentration, 20μL stock solution was mixed with 80 uL APEL-2. From this solution 5.6μl has been pipetted to 10 mL 5 NPEM. Cycloheximide was used at 10μM concentration as it has shown efficiency previously in different in vitro systems (45, 46). For Vehicle control a matching DMSO solvent control was used. NPCs were seeded at 300 000 cells/well and CHX treatment has been applied for 12 hours simultaneously with the 5μM CHIR induction. Cell viability has been evaluated via the assay of LIVE/DEAD™ Viability/Cytotoxicity Kit, for mammalian cells (ThermoFisher Scientific). Media was then aspirated off and cells were trypsinized. Enzymatic reaction was stopped with autoMACS running buffer + 10% FBS. After centrifuging at 3 min and 500rpm, we followed the LIVE/DEAD™ Viability/Cytotoxicity Kit, for mammalian cells (ThermoFisher Scientific) protocol. Images of cell suspension were acquired by Leica Thunder system with 10x 0.45 NA objective with green and red filter cubes on 24-well cell culture plates. Percentage of dead/live cells were quantified and calculated by Imaris using spot module.

### In vitro mRNA synthesis

Cre mRNA was created using pCAG-Cre plasmid (Addgene Catalog number #13775), mCherry mRNA was created using (Addgene pX330-U6-Chimeric_BB-CBh-hSpCas9 Cat# 42230) and GFPmRNA was made using pCAG-GFP (Addgene Cat# 11150).

DNA template for RNA synthesis was created using the forward and reverse primers listed below with GXL prime star PrimeSTAR® GXL DNA Polymerase (Takara Cat# R050A). mMESSAGE mMACHINE™ T7 ULTRA Transcription Kit (ThermoFisher Cat# AM1345) was used for in vitro mRNA synthesis from DNA template to create 5’CAP 3’Polyadenylated tailed transcripts. Synthesized mRNA was precipitated by lithium chloride and ran on a 1.5% agarose formaldehyde denaturing gel to validate proper size as well as tailing. See Table 2 primers for making DNA Templates.

### Cell transfection

mRNAs as well as sgRNAs were transfected to NPCs using Lipofectamine™ MessengerMAX™ Transfection Reagent (Thermofisher, Cat # LMRNA015). Per 24 well, 500 ng of total mRNA (per transcript type) were added along with sgRNAs at 1 μM concentration. NPCs were transfected according to manual instructions in OPTI-MEM. NPCs were incubated with OPTI-MEM for 3 hours (See Figure 2A).

### CRISPR mediated gene removal

sgRNA targeting GFP were purchased from ThermoFisher true guide gRNA sequence targeting exon 1 of GFP and designed using Invitrogen True guide Tool. sgRNAs targeting Ctnnb1 were designed using indephi to maximize frameshift potential (Shen, 2018 #410) and cross referenced with (Hodgkins, 2015 #411) to minimize off target effects. We custom ordered Alt-R CRIPSR-Cas9 sgRNA,2nmol sgRNA’s from IDT. We designed 4 guides, one targeting exon 2, 2 targeting exon3 and one targeting exon6 of B-catenin. See Table 2 for sgRNA sequences.

Lef/TCF factor targeting using sgRNAs utilized 48 h KO period. sgRNA’s were ordered from Synthego and designed using CRISPR Design Tool based on the following criteria: (1) targeting an early coding exon, (2) targeting a common exon between all the transcripts, (3) high activity gene cutting and (4) minimal off target effects. The highest ranked top four guides were tested using IF staining to measure protein loss as a metric of KO efficiency. Lyophilized guides were reconstituted at 100 pmol/ul in TE buffer and stored at -20 °C and gRNAs were used at 7.5 pmol/well for a 24-well plate. We used 1.875 pmol/guide/well in case of QKOs of Tcf7l1, Tcf7l2, Tcf7, Lef1. sgRNA’s were co-transfected cells with mCherry (500 ng/well mRNA) using Lipofectamine™ MessengerMAX™ Transfection Reagent (Thermofisher, Cat # LMRNA015).

### RNA isolation and RT-qPCR

RNA was isolated with Rneasy micro kit (Qiagen, #74004). RNA was also isolated using this kit to send for bulk RNA-seq. For RT-qPCR, RNA was reverse transcribed in cDNA with SuperScript IV VILO Master Mix with ezDNase Enzyme (cat #: 11766050). qPCR was performed with Luna Universal qPCR Master Mix Protocol (New England Biolab #M3003) on a ViiA 7 Real-Time PCR 96 System. See Table 2 for primer sequences.

### FACS sorting mRNA isolation

Prior to FACS sorting, NPCs were rinsed once with PBS, then incubated with trypsin for 5 min in the incubator at 37 degrees Celsius. Reaction was quenched with 10% FBS in AutoMACS buffer. NPCs were we pelleted and washed once with AutoMACS buffer before resuspending AutoMACS with in DAPI (dead cell dye DAPI (4’,6-Diamidino-2-Phenylindole, Dilactate, 422801, BioLegend) and DRAQ5 (DRAQ™ Live cell dye, NBP2-81125-50ul, NOVUS). 60,000 to 150,000 mCherry+ NPCs were sorted on BD SORP FACS Aria Iiu into RLT buffer with 1:100 beta-mercaptoethanol prior to RNA isolation.

### RNA-seq

Total RNA integrity was determined using Agilent Bioanalyzer or 4200 Tapestation. Library preparation was performed with 10 ng of total RNA with a Bioanalyzer RIN score greater than 8.0. ds-cDNA was prepared using the SMARTer Ultra Low RNA kit for Illumina Sequencing (Takara-Clontech) per manufacturer’s protocol. cDNA was fragmented using a Covaris E220 sonicator using peak incident power 18, duty factor 20%, cycles per burst 50 for 120 seconds. cDNA was blunt ended, had an A base added to the 3’ ends, and then had Illumina sequencing adapters ligated to the ends. Ligated fragments were then amplified for 12-15 cycles using primers incorporating unique dual index tags. Fragments were sequenced on an Illumina NovaSeq-6000 using paired end reads extending 150 bases.

Basecalls and demultiplexing were performed with Illumina’s bcl2fastq2 software. RNA-seq reads were then aligned to the combined mouse GRCm38 and human GRCh38 Ensembl release 76 primary assemblies with STAR version 2.5.1a1. Gene counts were derived from the number of uniquely aligned unambiguous reads by Subread:featureCount version 1.4.6-p52. Isoform expression of known Ensembl transcripts were estimated with Salmon version 0.8.23. Sequencing performance was assessed for the total number of aligned reads, total number of uniquely aligned reads, and features detected. The ribosomal fraction, known junction saturation, and read distribution over known gene models were quantified with RseQC version 2.6.24.

Normalized counts tables were ran through DeSEQ2 (Love et al., 2014) to create differential gene expression tables with Log2FC cut offs no less than 1 (no less than 0.5 Table S1.2) and p-adjusted values no greater than 0.05 can be found in tabs within Table S1. Significance of Cre and Cas9 intersected gene lists was calculated with hypergeometric function in R. Differential expression tables were passed through GO package clusterProfiler (Yu et al., 2012). Data was visualized using ggplot and complex heatmap functions in R (Gu, 2022). Benjamini-Hochberg correction (False Discovery Rate) applied as well as significance associated with intersections compared to all genes expressed with gene normalized counts greater than or equal to 10 using hypergeometric test.

### Immunofluorescence staining

For immunofluorescence staining NPCs were cultured on coverslips (Thermanox plastic coverslip, Thermo Fisher Scientific, Cat #174969), cell cultures were fixed with 4% PFA in PBS for 10 min, then washed with PBS twice before blocking in 1.5% SEA block (Thermo Fisher Scientific, 107452659) in TBST (0.1% Tween-20 in TBS). After blocking room temperature for 1 hour, coverslips were switched to primary antibody (diluted in blocking reagent) incubation in 4 °C overnight. After washing three times with TBST, switched to secondary antibody (diluted in blocking reagent) incubation 1 hour min in room temperature, shielded from light. This was followed by three washes with TBST and a final rinse in PBS. Cover slips were then flipped onto coverglass (VWR® Micro Cover Glasses, Rectangular no 1.5 22×60, VWR Cat # 48393-221) onto 15 µl of mounting media (Fluoromount-G™ Mounting Medium (25 ml) Thermo Fisher Scientific Cat# 00-4958-02). After drying overnight, coverglass with cover slips were taped with double sided tape onto superfrost micro slides 25×75×1mm, cases (VWR, Cat # 48311-703). Slides were kept away from light at 4 °C prior to confocal imaging. Find primary and secondary antibodies used in Table2.

### Edu chasing

NPCs were chased with 10 μM Edu diluted in DMSO for 1 hour prior to fixation with 4% PFA. To visualize Edu incorporation, we used Click-iT™ EdU Cell Proliferation Kit for Imaging, Alexa Fluor™ 647 dye (Thermo Fisher Scientific, Cat# C10340). NPCs were stained prior to Click-it reaction.

### RNA-Scope

To perform in situ floursecent RNA analysis we used RNA scope Multiplex Fluorescent Reagent Kit v2 (Advanced Cell Diagnostics, Inc.). We followed the commercially available RNA-scope protocol: the tissue on slides were treated with hydrogen peroxide and protease, then hybridized with probes for 2 hours at 40 °C using the HybEZ oven (Advanced Cell Diagnostics, Inc.). Probes were then amplified and detected with tyramide signal amplification fluorescence. The slides were incubated with 1 mg/ml Hoechst 33342 (Molecular Probes). The tissue was imaged at 63X oil immersion lens using the Leica SP8 confocal microscope. The catalog numbers of probes from Advanced Cell Diagnostics, Inc. used in this work are listed as follow: Ovol1 (845271-C1), Emx2 (319001-C3).

### Image acquisition

Image acquisition was performed using Leica SP8-X confocal fluorescence imaging system (Leica Microsystems, Germany) in 1024×1024 pixels with a 63X Leica oil immersion objective (NA 1.6).

### Image Quantifications

We quantified the confocal images of NPC experiments using Imaris microscopy image analysis software (version 9.9, Oxford Instruments).β-catenin KO quantification:

We quantified the membrane levels β-catenin protein in the KO experiments by Imaris as well. Background noise has been reduced by applying ‘Background Subtraction’ feature on the channel of examined protein. We manually created five membrane masks by the ‘surfaces’ module per image, measuring the fluorescent intensity of the shared membrane area between two adjacent transfected (KO) cells of the protein of interest, and calculated an average of these values. We chose to quantify membrane intensity only between two transfected cells to avoid the potential overestimation of protein levels: the signal of a non-transfected cell could overlap with the membrane of a KO cell. We also applied the same quantification method for adjacent non-transfected cells (the average of five membrane intensities between non-transfected cells). We calculated a ratio between the average fluorescent membrane intensity between two transfected cells and the average membrane intensity of non-transfected cells. This ratio has been multiplied by 100 and reported as the percentage KO of the protein. The latter normalization step was required to correct the differences in immunostaining and image acquisition across samples.

Jag1 loss from β-catenin KO quantification: Cells were manually classified as Jag1+ if the nuclear masks had granular Jag1 signal in their proximity and/or the signal was observed in membrane distribution.

Edu quantification:Cells were classified as mCherry+ or Edu+ with the use of spots module based on cut-offs of their respective fluorescent intensity histograms in.

Jag1 staining and inside outside quantification time course β-catenin KO quantification: ”Spots” module was used to determine the number of mCherry+ cells outside the aggregates. Cells not in contact with Jag1+ aggregates were categorized as “outside the aggregate” cells.

Six2 protein/ Tcf7l1 protein level quantification in β-catenin KO: Six2 and Tcf7l1 intensity was determined based on the nuclear intensity values of cell module.

Tcf/Lef KO and loss of Jag1 protein with QKO quantification: The number of cells were automatically counted by the software using the DAPI nuclear marker after setting the appropriate nuclear radius. The cells were then classified as mCherry+ based on manual cut-offs of their respective fluorescent intensity histogram. These mCherry+ cells were further classified as Lef1+ based on manual cut-offs of their respective fluorescent intensity histogram. We calculated the decrease of protein levels of Lef1 by finding the ratio between the Lef1+ mCherry+ cells and total mCherry+ cells. This ratio has been multiplied by 100 and reported as the percentage KO of the protein. This quantification process was also applied to determine the decrease of protein levels of Tcf7, Tcf7l1, Tcf7l2, and Jag1.

### Statistics

Initially, the normal distribution of datasets was determined by D’Agostino-Pearson test. In case of normal distribution, the p-values were calculated by Student’s unpaired t-test. Mann-Whitney test was applied for datasets with non-normal distribution.p<0.05 was considered significant. We quantified the individual and quadruple knockout (KO) experiments using Imaris microscopy image analysis software (version 9.9, Oxford Instruments).

### Integration of scRNA-seq (mouse and human) and ChIP seq data with bulk RNA seq Data

Previously published β-catenin and Tcf/Lef transcription factor ChIP-seq data (16) was downloaded from publicly available data depository (GEO accession: GSE131117). Raw sequencing reads of input, Ctnnb1, Lef1, Tcf7 ChiP-seq from low/high-concentration (1.25uM, 5uM) CHIR-treated NPCs were aligned by Bowtie2 (47) on mm39 reference genome. Alignment files were sorted by samtools (Li et al., 2009)and filtered to remove duplicate reads by picard ‘markduplicates’ tool (http://broadinstitute.github.io/picard/). Peak calling was performed by MACS2 with a q-value cutoff threshold of 10-8, to ensure strong transcription factor bindings in high-concentration CHIR treatment compared to low concentration counterparts. Peak annotation was performed by annotatePeaks.pl from HOMER suite (Heinz et al., 2010) with default parameter. To incorporate any β -catenin/TCF/LEF downstream genes in high CHIR condition, the union set of annotated genes were used in the following intersection analysis. Down-regulated DEGs in both Cas9- and Cre-mediated knockout bulk RNA-seq data were intersected with ChIP-seq peak associated genes. Lastly, 161 genes in the intersection list were visualized with Seurat (Stuart et al., 2019). FeaturePlot function in mouse post-neonatal day0 (P0) nephrogenic-lineage cells from kidney single-cell data (Kim et al., 2023 preprint). Total number of expressed genes in nephrogenic cells were calculated based on the threshold of more than 0.25% cells with normalized expression value of a gene.

### Human scRNA-seq analysis

To investigate conservation between mouse and human gene expression the 161 genes in the intersection mouse ChIP list were visualized and homologs were intersected with Seurat FeaturePlot function in human fetal kidney (Lindström et al., 2021).

## Acknowledgments

We thank former and current members of the McMahon laboratory for helpful comments, technical assistance and useful discussion especially Jill McMahon, JinJin Guo, Muskaan Singh, Dr. Tracy Tran, Dr. Kevin Peterson and Dr. Qiuyu Guo. We thank Dr. Christopher Garcia and Dr. Yi Miao for sharing the Wnt mimetic bi-specific antibody. We are grateful for the assistance of Dr. Seth Ruffins in the biological imaging core facility and Bernadette Masinsin in flow core facility and thank our colleagues Dr. Yulia Schwartz and Dr. Unmesh Jadhav for illuminating discussions.

## Author contributions

HB and APM conceived the study. HB and BD, performed experiments and collected data. HB, BD, NOL and SK performed various analyses on the data. HB and APM wrote the manuscript incorporating comments from all authors.

## Data deposition

The bulk mRNA-seq for this study can be found at GEO accession number (GSE231753). The mouse embryonic kidney single cell RNA is deposited at GEO accession number (GSE232482) the human fetal kidney single data is deposited at at GEO accession number (GSE139280). The GEO accession number for the bulk mRNA seq data in the corresponding manuscript is (GSE232606).

## Diversity and inclusion statement

In order to advance our educational, scientific and clinical mission, USC Stem Cell is deeply committed to creating a culture that supports diversity, equity and inclusion (DEI). HB identifies as an underrepresented minority female in STEM and benefitted from funds from the Ruth L. Kirschstein National Research Service Award (NRSA) Individual Predoctoral Fellowship to Promote Diversity in Health-Related Research (Parent F31-Diversity) and F31 DK122777.

## Figures Legends

**Figure S1:**
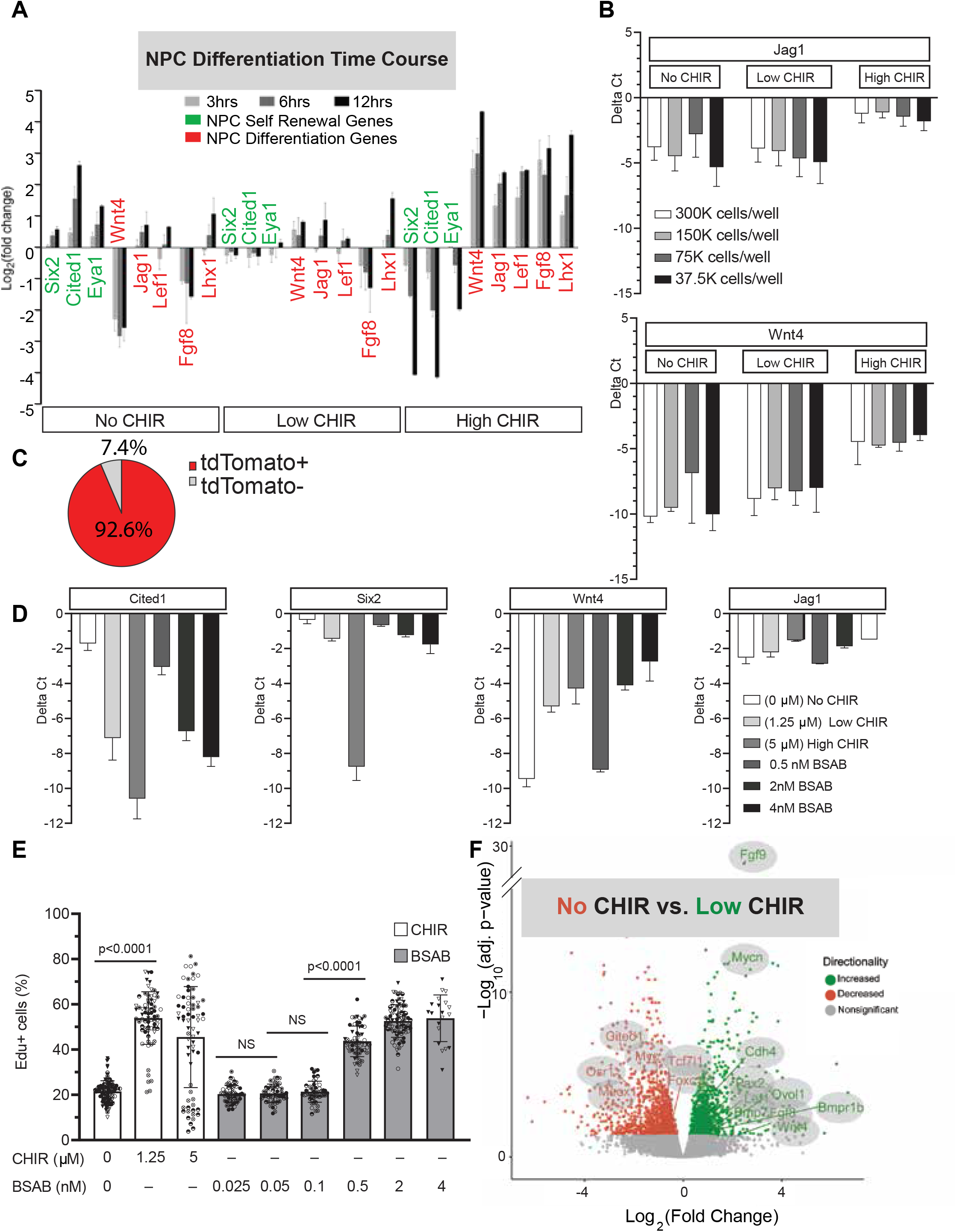
A) RT-qPCR of NPC cultured across a 3-, 6- and 12-hour time course. Values are normalized to Gapdh and a ratio is calculated by dividing low CHIR condition values. Values are Log2 transformed. Green genes are self-renewing NPC markers and red genes are NPC differentiation markers.. Grey panel denotes no (0μm) CHIR, light gray denotes low (1.25 μm) CHIR and dark gray denotes high (5μm) CHIR after a 24 hour low CHIR seeding stabilization period (Biological replicates n= 3). B) RT-qPCR results for Six2, Cited1, Wnt4, Lef1, Lhx1, Jag1 (Gapdh-Ct value) of NPC cultured at 300k, 150k, 75k and 37.5k at 0 μm, 1.25 μm and 5 μm CHIR. Values placed further from x-axis denote less expression. Delta Ct = (Gapdh – Ct gene value) (Biological replicates n= 3). C) Percent purity of NPCs derived from Six2TGC-tdTomato mice. Purity measured as cell numbers of cells that are tdTomato + as a result of Six2-Cre activity (Range from 86%-100% from 32 field of view from 8 wells. D) RT-qPCR of self-renewal genes (Cited1, Six2) and induction genes (Wnt4, Jag1) of NPCs cultured with Wnt surrogate BSAB to validate the activation of Wnt pathway using CHIR. n= 2 biological replicates. ΔCt = (Gapdh – Ct gene value). E) Quantification of Edu chasing (1 hour) of NPCs post 24-hour exposure to NPEM with different types and various levels of Wnt input (CHIR and BSAB, n=2-3 biological replicates), Mann-Whitney test, unpaired t test. F) Bulk RNA-seq data of DEGs using Log2FC = absolute value cut off=0.5 and p-adjusted value cut off =0.05 comparing no (0μm) CHIR and low (1.25μm) CHIR post 24 hours of culture represented as a volcano plot.

**Figure S2:**
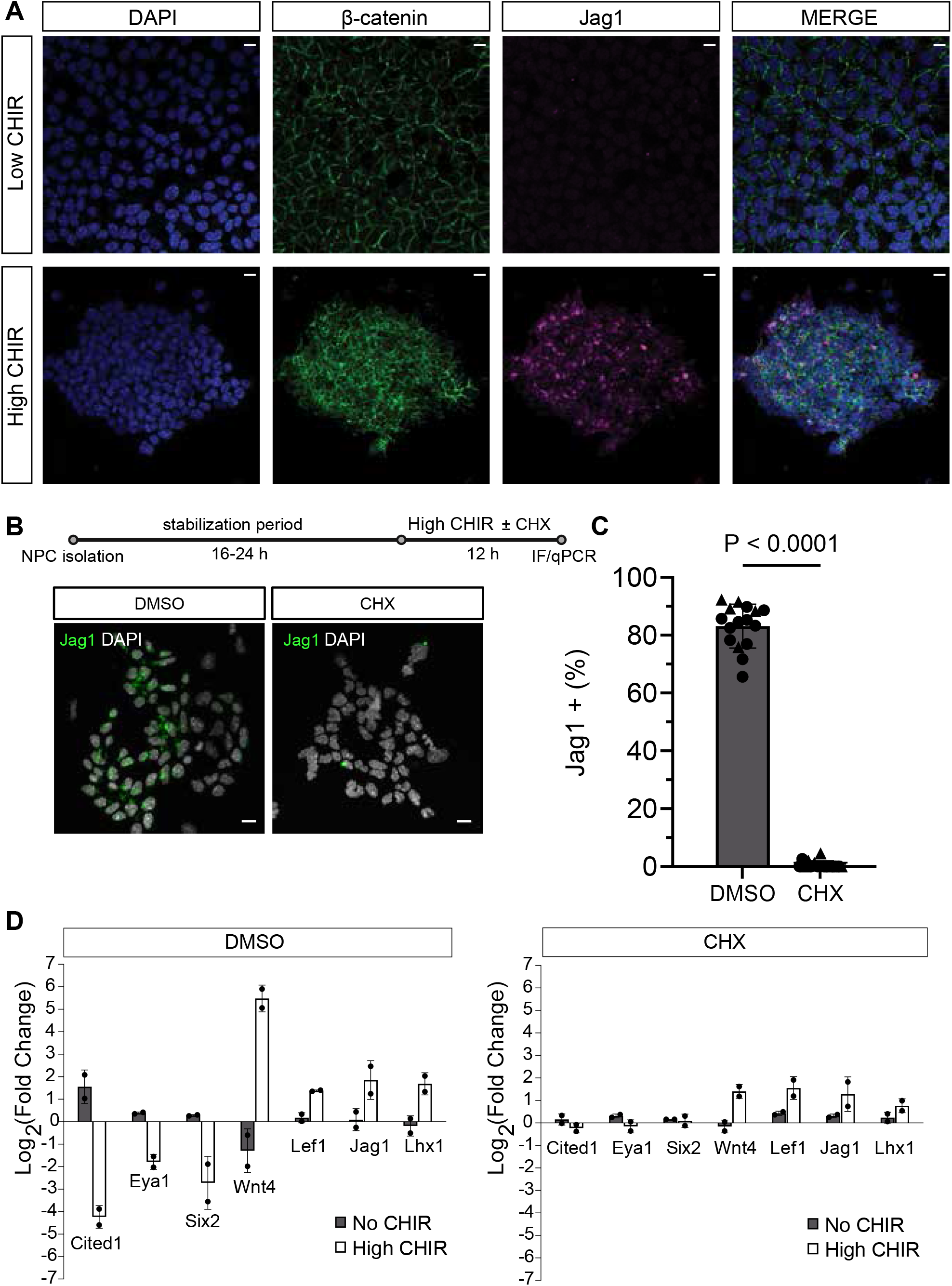
A) Immunofluorescent staining of β-catenin (green), Jag1 (purple), DAPI (blue) on wildtype NPC cultured in low (1.25 μM) and high (5 μm) CHIR (scale bar = 10 μm). B) Immunofluorescent staining of NPCs treated with CHX for 12 hours (scale bar = 10 μm). F) Quantification of Ja1 expression in CHX treated NPCs vs. vehicle (DMSO) control. n=2 biological replicates denoted as different symbols, 16 field of views/group. Wilcoxon-test. E) RT-qCPR of NPCs treated with CHX for 12 hours. Log2FC graphed compared to low CHIR conditions. Values normalized to Gapdh expression.

**Figure S3:**
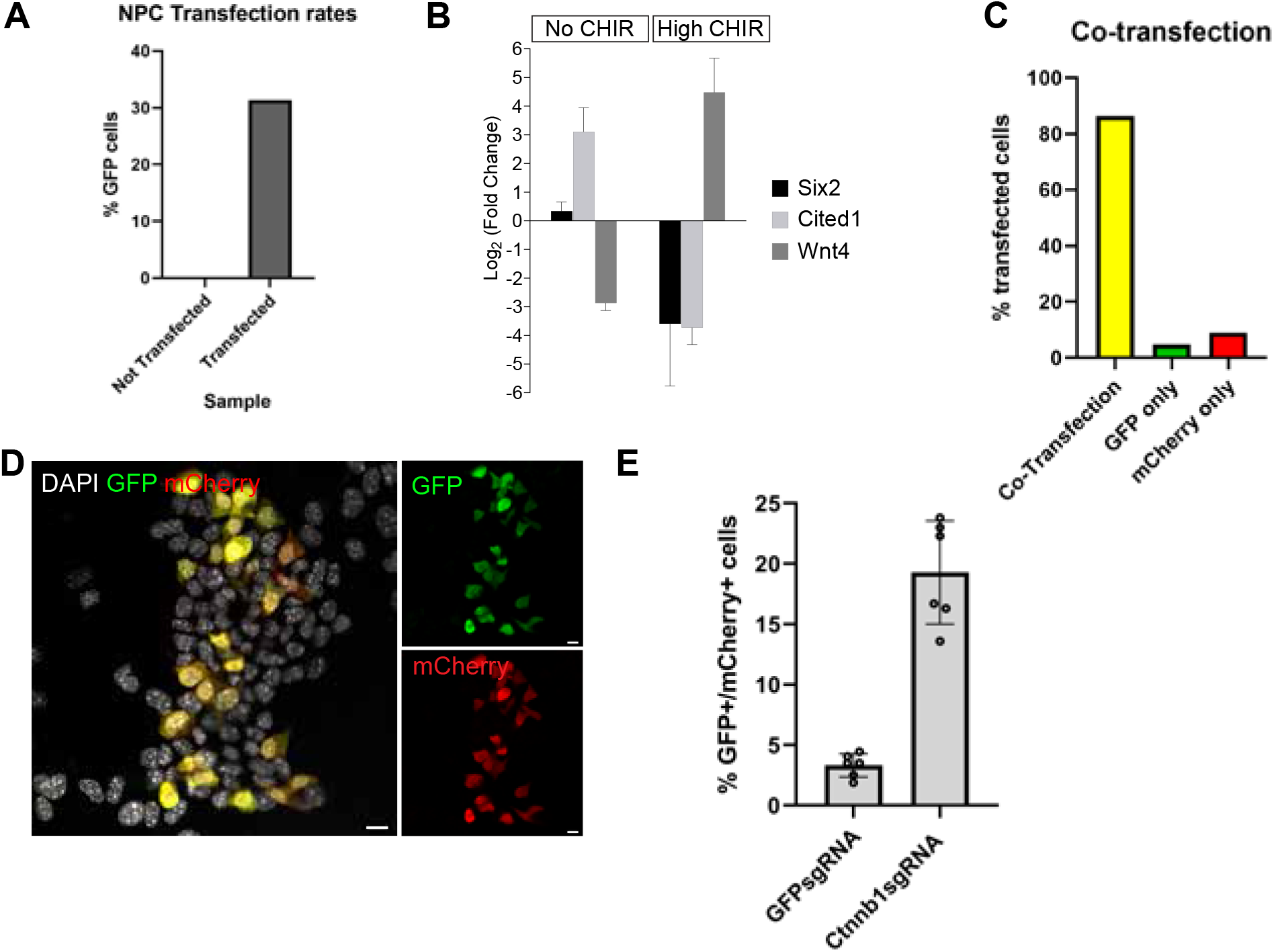
A) Transfection rates of GFP-mRNA in NPCs measured by FACS sorting (n=2 biological replicates) B) RT-qPCR of NPCs transfected with GFP and harvested 24 hours after changing media with altering CHIR conditions. Ct-differences normalized to GAPDH and samples compared to 1.25 μm CHIR condition. Log2 transformed values are plotted. (n=2 biological replicates). C) FACS sorting measuring co-transfection (mCherry mRNA and GFP mRNA) rates of mCherry and GFP mRNA into NPCs transfected together (co-transfection) or independently (n=2 biological replicates). D) Immunostaining of GFP and mCherry in NPCs demonstrating co-localization of GFP (green) and mCherry (red) appearing yellow (green + red) (scale bar = 10 μm). E) GFP positive NPCs in GFP sgRNA/mCherry co-transfected samples and Ctnnb1 sgRNA/mCherry co-transfected samples. GFP signal reduction as a result of GFP sgRNA controls.

**FigureS4 :**
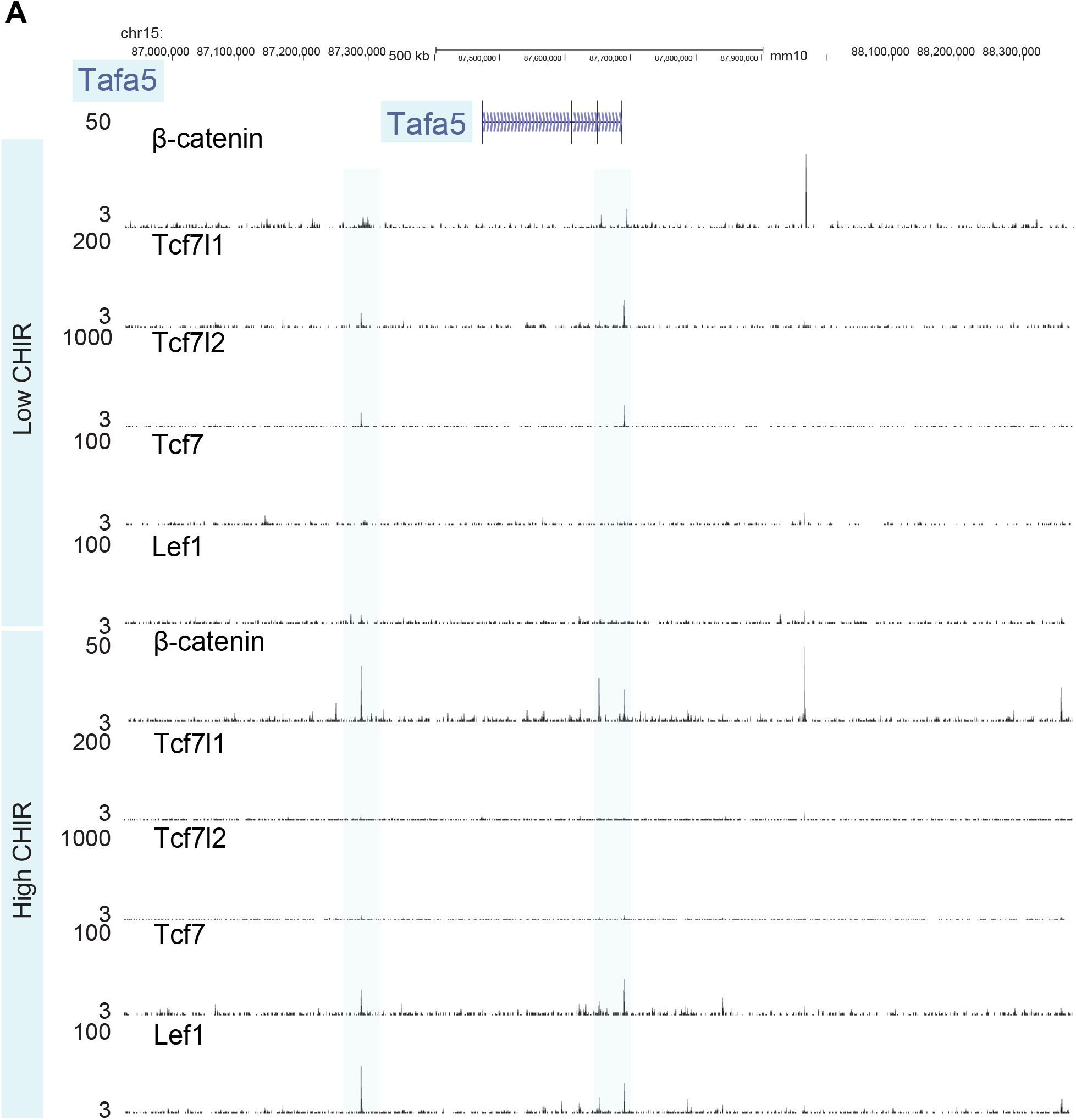
A)) Ctnnb1/Tcf7l1/Tcf7l2/Tcf7/Lef1 ChiP seq at Tafa5 (Fam19a5) cis-regulatory regions express induction marker Tcf/Lef switch previously reported in induction target genes.

**Figure S5:**
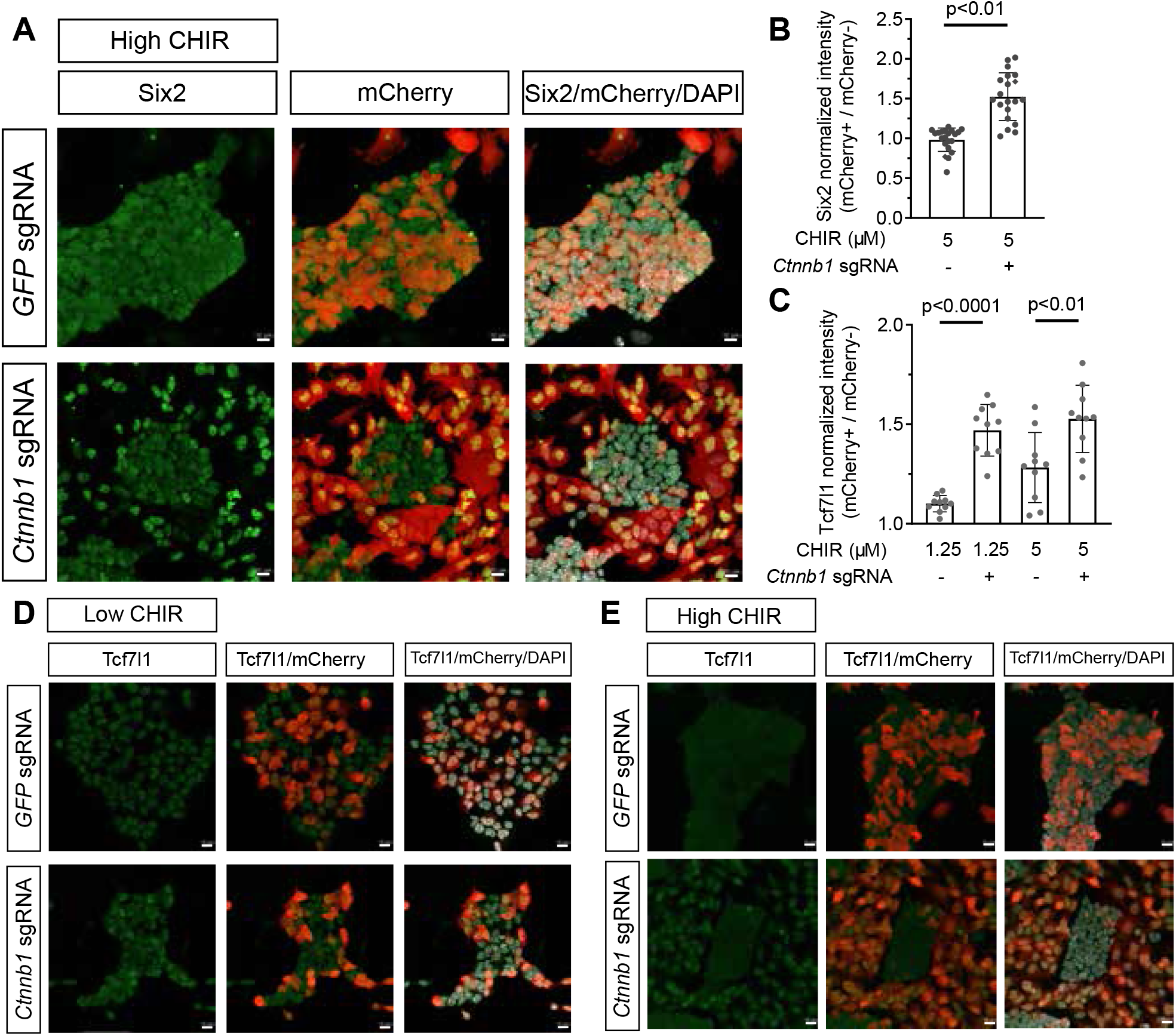
A) Immunostaining of NPCs with Cas9 mediated β-catenin KO in high CHIR. B) Quantification of the increase in Six2 immunofluorescent intensity in Cas9 mediated β-catenin removal of NPCs cultured in high CHIR. Unpaired t test. 4-8 fields of view,3 biol repl, normality test pass, Student t test C) Quantification of increase in Tcf7l1 staining in Cre mediated β-catenin removal of NPCs Unpaired t test. 10 fields of view,1 technical replicate, normality test pass, Student t-test. D) Immunostaining of NPCs cultured in 1.25 uM CHIR. Tcf7l1= green, mCherry = Red. E) Immunostaining of NPCs cultured in 5uM CHIR. Tcf7l1= green, mCherry = Red

**Figure S6:**
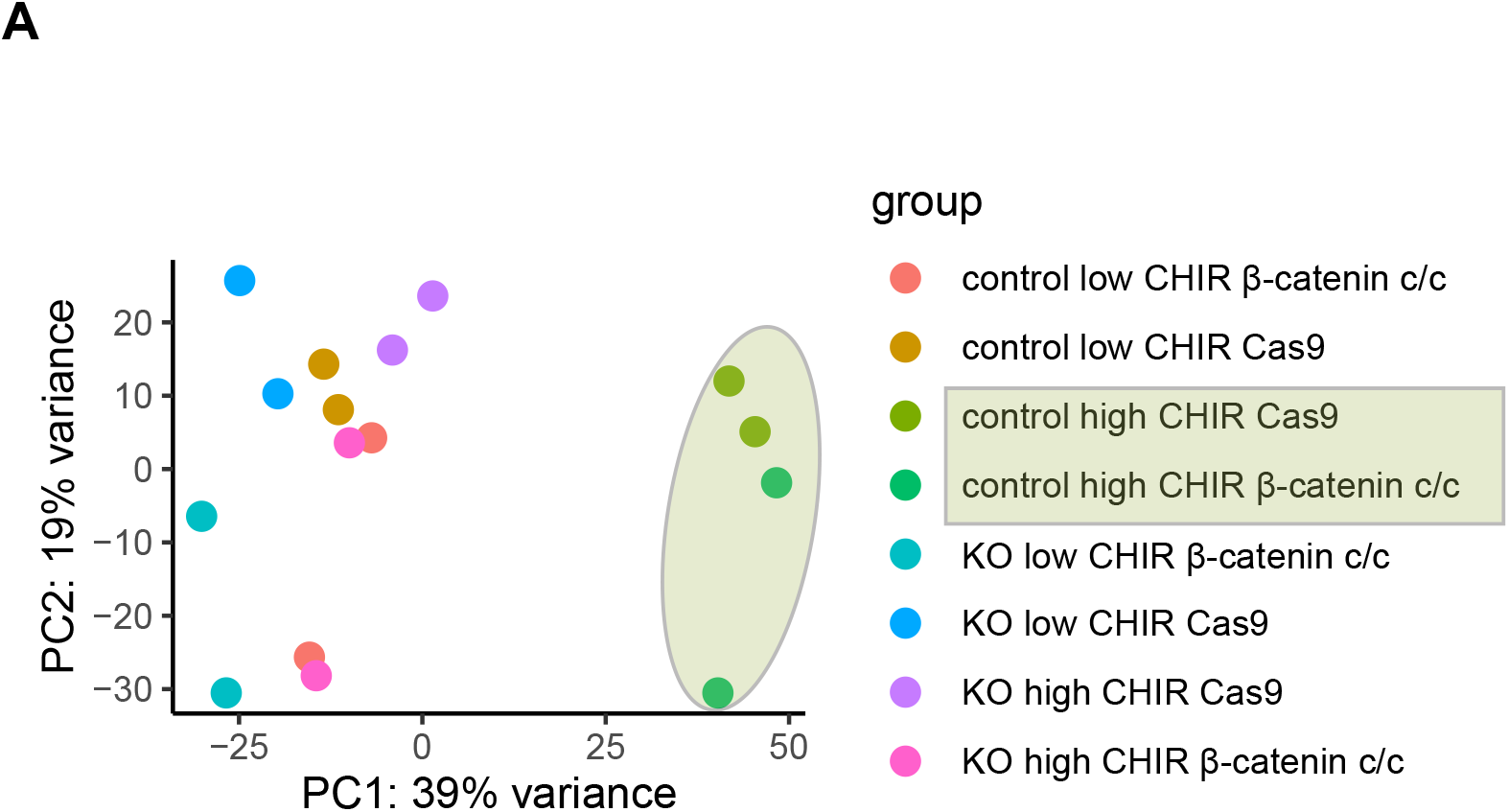
A) PCA plot of bulk mRNA-seg β -catenin KO samples.

**Figure S7:**
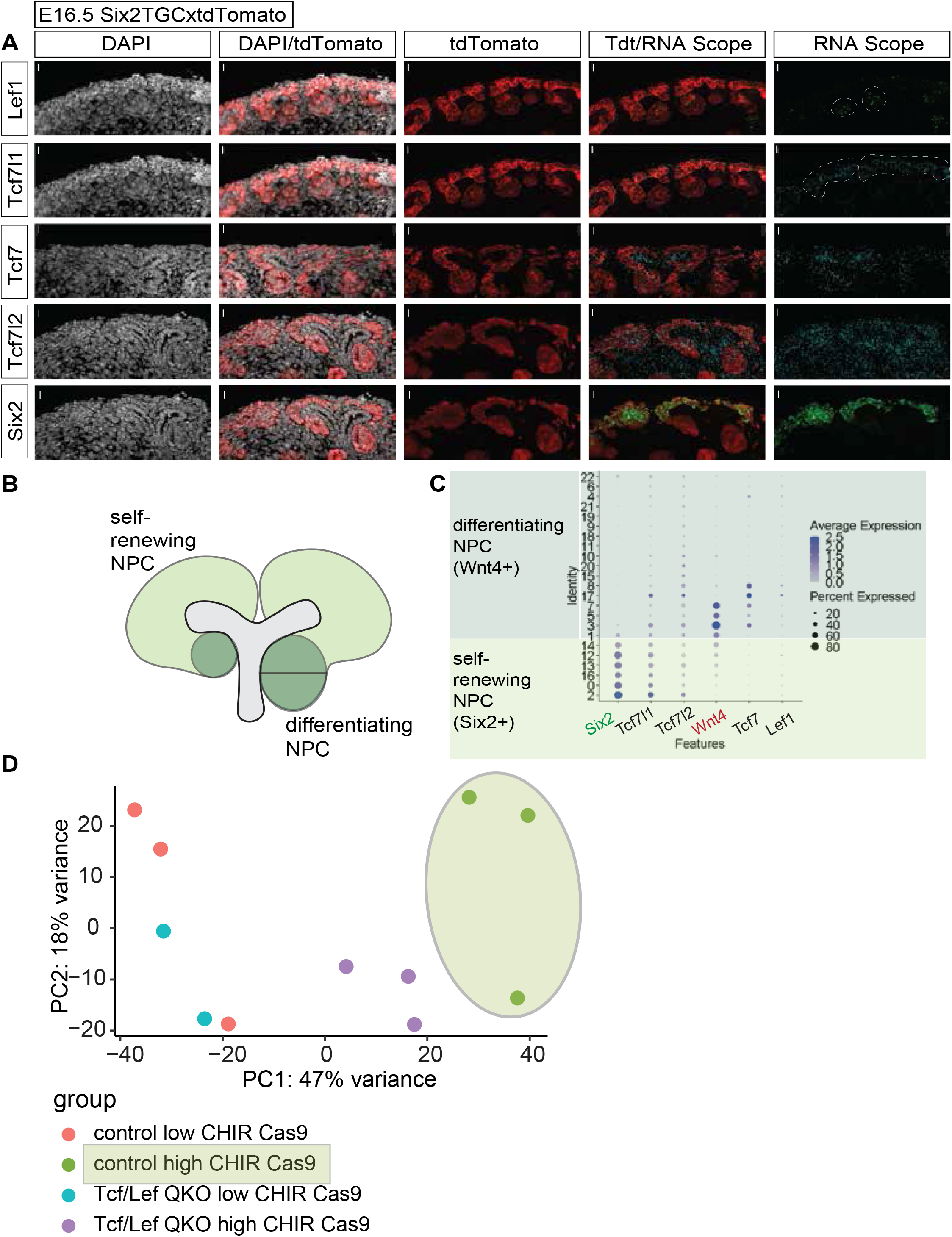
A) RNA scope of E16.5 Six2TGC x Tdt kidneys. Cyan = Tcf7l1, Lef1= Green, tdT antibody = red, DAPI= white. B) RNA scope of E16.5 Six2TGC x Tdt kidneys. Cyan = Tcf7l2, DAPI= white, tdT antibody = red C) RNA scope of E16.5 Six2TGC x Tdt kidneys. Cyan = Tcf7, Lef1 antibody = Green, tdT antibody = red, DAPI= white. D) scRNA-seq E16.5 kidneys – (16) dotplot showing Tcf7l1 and Lef1 expression in self-renewing and induced NPCs respectively. Anchor genes Six (green) denotes self-renewing NPCs and Wnt4 (red) denotes induced NPCs. E) PCA plot of control (GFP sgRNA) and QKO (Tcf7l1, Tcf7l2, Tcf7, Lef1 sgRNA) in low and high CHIR

**Figure S8:**
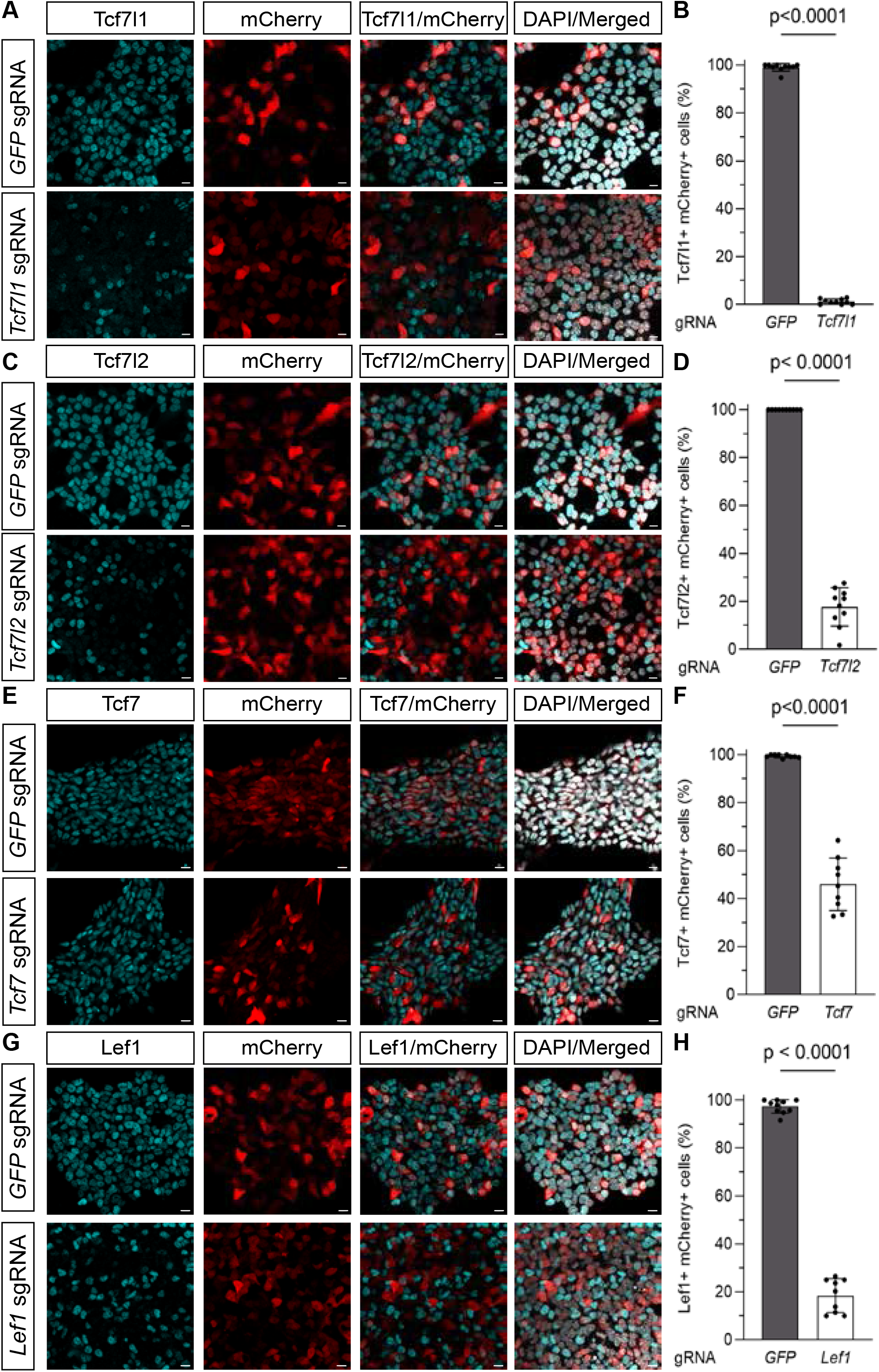
A) Immunofluorescence staining denoting Tcf7l1 protein removal (10μm scale bar). B) Quantification of Tcf7l1 protein removal. Mann-Whitney test, 10 field of views/well. C) Immunofluorescence staining denoting Tcf7l2 protein removal (10μm scale bar). D) Quantification of Tcf7l2 protein removal. 10 field of views/well. Unpaired t test. E) Immunofluorescence staining denoting Tcf7 protein removal (10μm scale bar). F) Quantification of Tcf7 protein removal., 9 field of views/well. Unpaired t test. G) Immunofluorescence staining denoting Lef1 protein removal (10μm scale bar). H) Quantification of Lef1 protein removal. 9-10 field of views/well. Unpaired t test.

**Figure S9:**
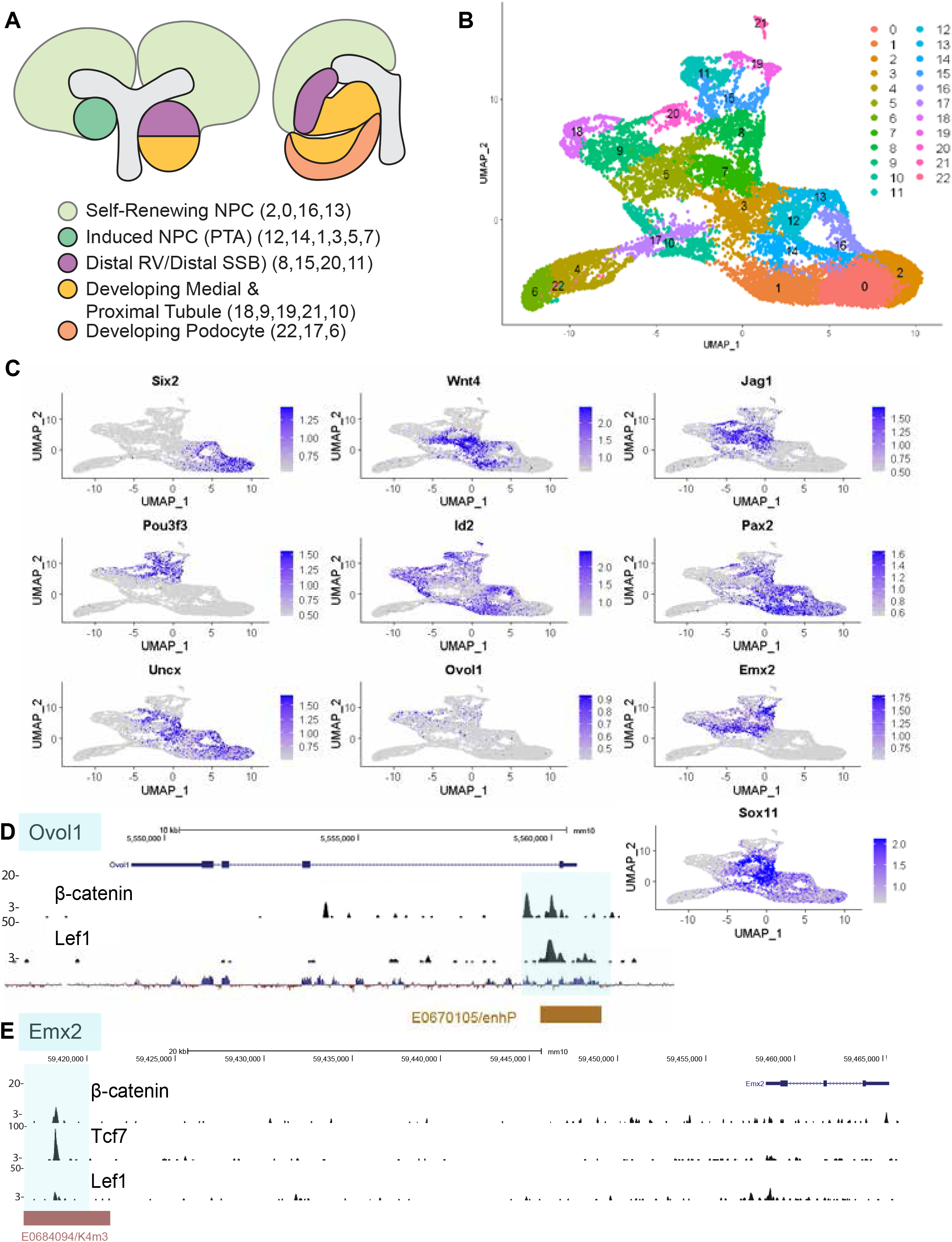
A) Schematic representation of mouse nephrogenic niche and developing s-shape body B) UMAP of all clusters of nephrogenic linage of cells from p0 scRNA-seq data (Kim et al., 2023 preprint) C) feature plots of β-catenin target genes Pax2, Id2, Uncx, Emx2, Ovol1, Dach1, Sox11 with known markers of self-renewing NPCs (Six2, Cited1), early induction gene (Wnt4), Distal marker (Pou3f3), proximal tubule marker (Hnf4a), podocyte gene (Mafb) in mouse p0 scRNA-seq D) gene tracks of ChiP-seq data at Ovol1 locus showing β-catenin/ Lef1 binding at annotated cis-regulatory module of Ovol1 gene E) gene tracks of ChiP-seq data at Emx2 locus showing β-catenin/ Lef1/Tcf7 binding at annotated cis-regulatory module of Emx2 gene

